# PLD3 and PLD4 synthesize *S,S*-BMP, a key phospholipid enabling lipid degradation in lysosomes

**DOI:** 10.1101/2024.03.21.586175

**Authors:** Shubham Singh, Ulrich Dransfeld, Yohannes Ambaw, Joshua Lopez-Scarim, Robert V. Farese, Tobias C. Walther

## Abstract

Bis(monoacylglycero)phosphate (BMP) is an abundant lysosomal phospholipid required for degradation of lipids, in particular gangliosides. Alterations in BMP levels are associated with neurodegenerative diseases. Unlike typical glycerophospholipids, lysosomal BMP has two chiral glycerol carbons in the *S* (rather than the *R*) stereo-conformation, protecting it from lysosomal degradation. How this unusual and yet crucial *S,S-*stereochemistry is achieved is unknown. Here we report that phospholipases D3 and D4 (PLD3 and PLD4) synthesize lysosomal *S,S-*BMP, with either enzyme catalyzing the critical glycerol stereo-inversion reaction *in vitro*. Deletion of PLD3 or PLD4 markedly reduced BMP levels in cells or in murine tissues where either enzyme is highly expressed (brain for PLD3; spleen for PLD4), leading to gangliosidosis and lysosomal abnormalities. PLD3 mutants associated with neurodegenerative diseases, including Alzheimer’s disease risk, diminished PLD3 catalytic activity. We conclude that PLD3/4 enzymes synthesize lysosomal *S,S-*BMP, a crucial lipid for maintaining brain health.

## INTRODUCTION

The degradation of lipids (for instance from lipoproteins, lipid droplets, or cell membranes) by acid hydrolases is a major function of lysosomes. Lysosomal lipid degradation is thought to occur predominantly in the lysosomal lumen, because the delimiting membrane of lysosomes is protected from acid hydrolases by an extensive glycocalyx. The degradation of membrane lipids in the lysosomal lumen requires the formation of intralumenal vesicles (ILVs) to provide acid hydrolases access to membrane lipids. ILV membranes differ in composition from other cell membranes. They are enriched with a specific phospholipid, bis(monoacylglycero)phosphate (BMP), which constitutes up to 70 mol% of ILV phospholipids (Kobayashi et al., 2002) and may be involved in ILV formation (Frederick et al., 2009; Matsuo et al., 2004). A key feature of BMP is that it is negatively charged at lysosomal pH (4.5–5.0), which is thought to enable binding of positively charged regions of specific acid hydrolases to ILV membranes (Anheuser et al., 2015; Gruenberg, 2020; Oninla et al., 2014; Sandhoff and Sandhoff, 2018). BMP-mediated lipid degradation is crucial for cells. For example, BMP deficiency leads to defects in the degradation of ganglioside lipids (sialic acid–containing glycosphingolipids that are abundant in the brain) (Svennerholm, 1980), and excessive ganglioside accumulation appears to be toxic to the nervous system (Breiden and Sandhoff, 2019; Schulze and Sandhoff, 2014). Additionally, alterations in BMP-mediated lysosomal lipid degradation are increasingly recognized as a hallmark of neurodevelopmental and neurodegenerative diseases (Boland et al., 2022; Logan et al., 2021; Udayar et al., 2022).

To maintain BMP levels in ILVs, BMP must avoid degradation by lysosomal phospholipases. How does this lipid resist lysosomal degradation? This property is apparently provided by the unique stereochemistry of lysosomal BMP. BMP has two glycerol molecules, and lysosomal BMP has both glycerol chiral carbons in the *S-* rather than the *R-*stereo-conformation that is typically found in glycerophospholipids (**Figure S1A**) (Amidon et al., 1995; Brotherus et al., 1974; Tan et al., 2012b; Taniguchi et al., 2015). The *S,S-*configuration is thought to protect BMP from degradation by lysosomal acid phospholipases (Abe and Shayman, 2009; Grabner et al., 2019; Gruenberg, 2020; Showalter et al., 2020).

The key question for understanding BMP synthesis is therefore how cells synthesize the unique *S,S*-stereo-isoform of BMP. BMP is thought to be synthesized in a multistep pathway from the precursor *R,S*-phosphatidylglycerol (PG) (Haverkate and Van, 1964; Hostetler, 1982) (**Figure 1A**). Inasmuch as the precursor lipid PG is an isomer of BMP, its synthesis pathway can be viewed as requiring two principal steps: a rearrangement of acyl moieties around the glycerol-phosphate-glycerol backbone, and a stereo-conversion of a *R*-glycerol to *S*-glycerol to generate a molecule with both glycerols in the *S*-configuration. The latter step is crucial to yield *S,S-*BMP, which is the form found in mammalian lysosomes (Amidon et al., 1995; Brotherus et al., 1974; Joutti, 1979; Tan et al., 2012b; Taniguchi et al., 2015).

**Figure 1.**
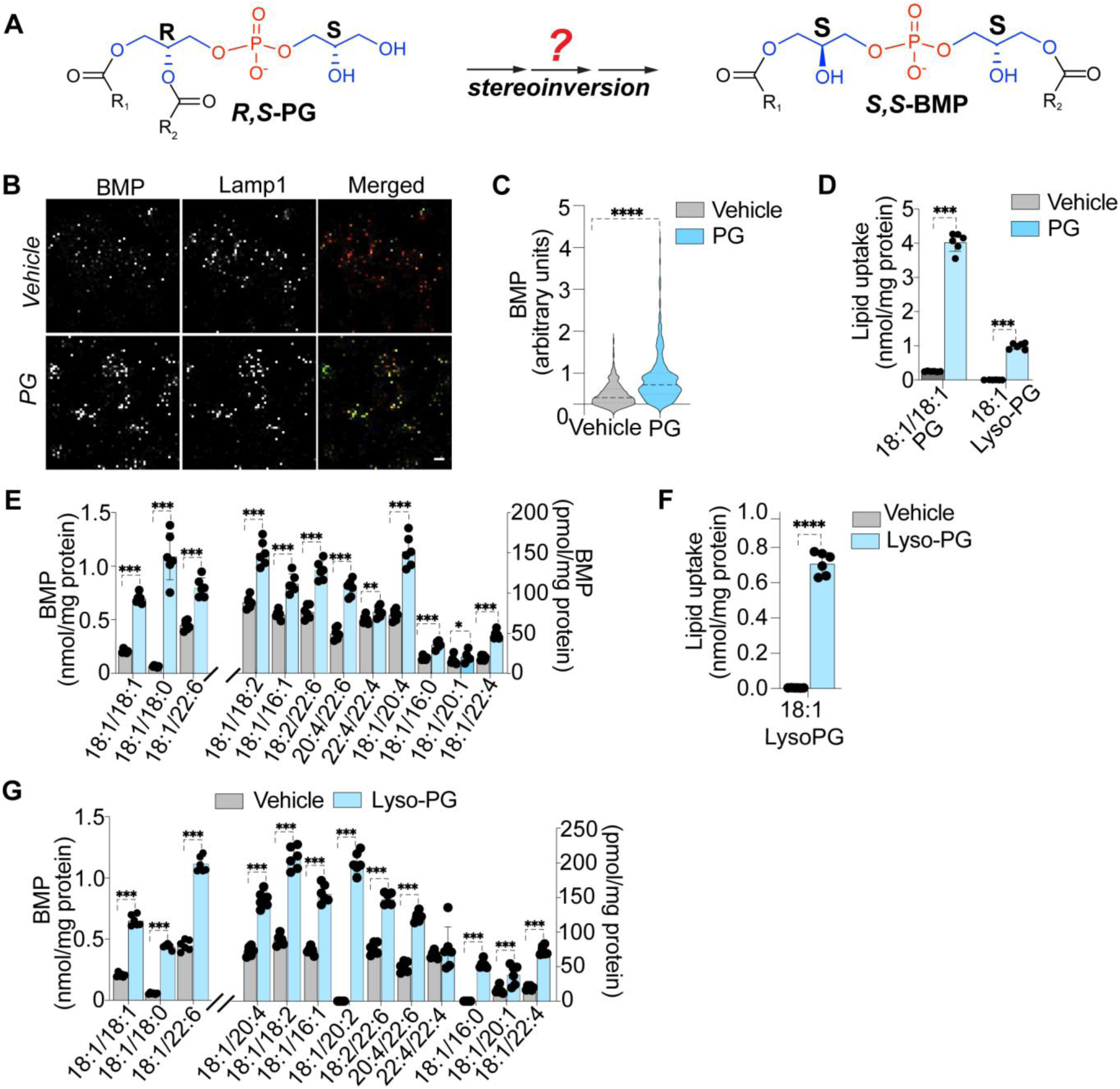
PG and lyso-PG are precursors of BMP in mammalian cells. (A) Scheme of BMP synthesis involves an unclear but crucial stereo-inversion step to yield *S,S*-BMP from *R,S*-PG. The two glycerol molecules are shown in blue. (B) Confocal microscopy images of anti-BMP (green) and anti-LAMP1 (red) staining of HMC3 cells show that PG feeding increased BMP levels in the lysosomes. Scale bar =10 μm. (C) Quantification of BMP signal (green) over masked LAMP1 (red) signal demonstrates ∼two fold increases in BMP levels. \*\*\*\**p <* 0.0001 vehicle vs. lipid feeding (Student’s t-test). (D) Incubation of HMC3 cells with 18:1/18:1 PG for 6 h increased 18:1/18:1 PG and 18:1 lyso-PG levels. (E) Incubation of HMC3 cells with 18:1/18:1 PG for 6 h increased levels of BMP. (F) Incubation of HMC3 cells with 18:1 lyso-PG for 6 h increased 18:1 lyso-PG levels. (G) Incubation of HMC3 cells with 18:1 lyso-PG for 6 h increased BMP species. PG and lyso-PG used in the assays were in *R,racemic* form. Data are mean ± S.D. (n=6 per group). \**p <* 0.05; \*\**p <* 0.01; \*\*\**p <* 0.001 vehicle vs. lipid feeding (multiple t-test).

Here, we report the identification of the lysosomal PLD3 and PLD4 as enzymes that mediate the synthesis of *S,S-*BMP. We show that these enzymes, expressed in cultured cells or as purified enzymes, catalyze BMP synthesis, and that the product of these enzymes is lysosomally stable *S,S*-BMP. Further, these enzymes are required to maintain BMP levels in cells or tissues where they are expressed, and the loss of these enzymes leads to BMP deficiency, ganglioside accumulation, and lysosomal abnormalities.

## RESULTS

### Human cells synthesize BMP from PG or lyso-PG

Prior studies reported that BMP may be synthesized from PG (Amidon et al., 1995; Hullin-Matsuda et al., 2007; Somerharju and Renkonen, 1980; Waite et al., 1987). We thus tested this hypothesis in human microglia clone 3 (HMC3) cells. Microglial cells were chosen as being particularly relevant to ganglioside degradation in the central nervous system. Incubation of cells with medium containing 18:1/18:1 PG resulted in a ∼twofold increase in cellular BMP levels, as assessed by immunofluorescence microscopy (**Figure 1B,C**). To independently measure changes in BMP under these conditions, we developed and validated a chromatographic separation and LC-MS/MS mass spectrometry protocol that reliably distinguished between PG and BMP lipid species, which are isobaric and thus often incorrectly measured together (**Figure S1B-G**). As expected, the analysis of cellular lipids by LC-MS/MS revealed a ∼10-fold increase of 18:1/18:1 PG and a ∼20-fold increase of lyso-PG in response to 18:1/18:1 PG incubation (**Figure 1D**). Additionally, total BMP levels increased by two- to threefold, and analysis of BMP species revealed that numerous 18:1-containing BMP species were increased (**Figure 1E**). We also detected accumulation of PG species with different fatty acyl moieties, suggesting that re-acylation of 18:1 lyso-PG to PG occurred during the assays (**Figure S2A**). The same experiments performed in another human cell line (HEK293T) yielded similar results (**Figure S2B,C**).

Our data suggested that BMP is synthesized from lyso-PG that is generated by PG hydrolysis. Indeed, incubation of either HMC3 and HEK293T cells with medium containing 18:1 lyso-PG increased lyso-PG levels and resulted in ∼2–3-fold increases of different BMP species (**Figures 1F,G and S2D,E**). In contrast, incubation of cells with a variety of other glycerophospholipids did not increase cellular BMP levels, and instead increased levels of lysophospholipid derivatives (**Figure S2F-K**). Taken together, these data indicate that lyso-PG is a precursor of cellular BMP in mammalian cells.

### Human cells synthesize BMP by transphosphatidylation of lyso-PG in lysosomes

A key aspect of BMP synthesis is the generation of the lysosomally stable *S,S*-form of BMP. We therefore sought to verify that *S,S*-BMP is more stable in purified lysosomes than other stereo-isoforms of BMP, as proposed (Abe and Shayman, 2009; Grabner et al., 2019; Joutti, 1979; Shayman et al., 2011; Showalter et al., 2020). To test this, we incubated *R,R-*, *R,S-*, or *S,S*-stereoisomers of 18:1/18:1 BMP with lysosomal extracts from HMC3 cells. Indeed, both the *R,R-* and *R,S-* stereoisomers of BMP were degraded to lyso-PG in the lysosomal extracts of HMC3 cells, whereas *S,S-*BMP was much more resistant to degradation (**Figures 2A and S2L**).

**Figure 2.**
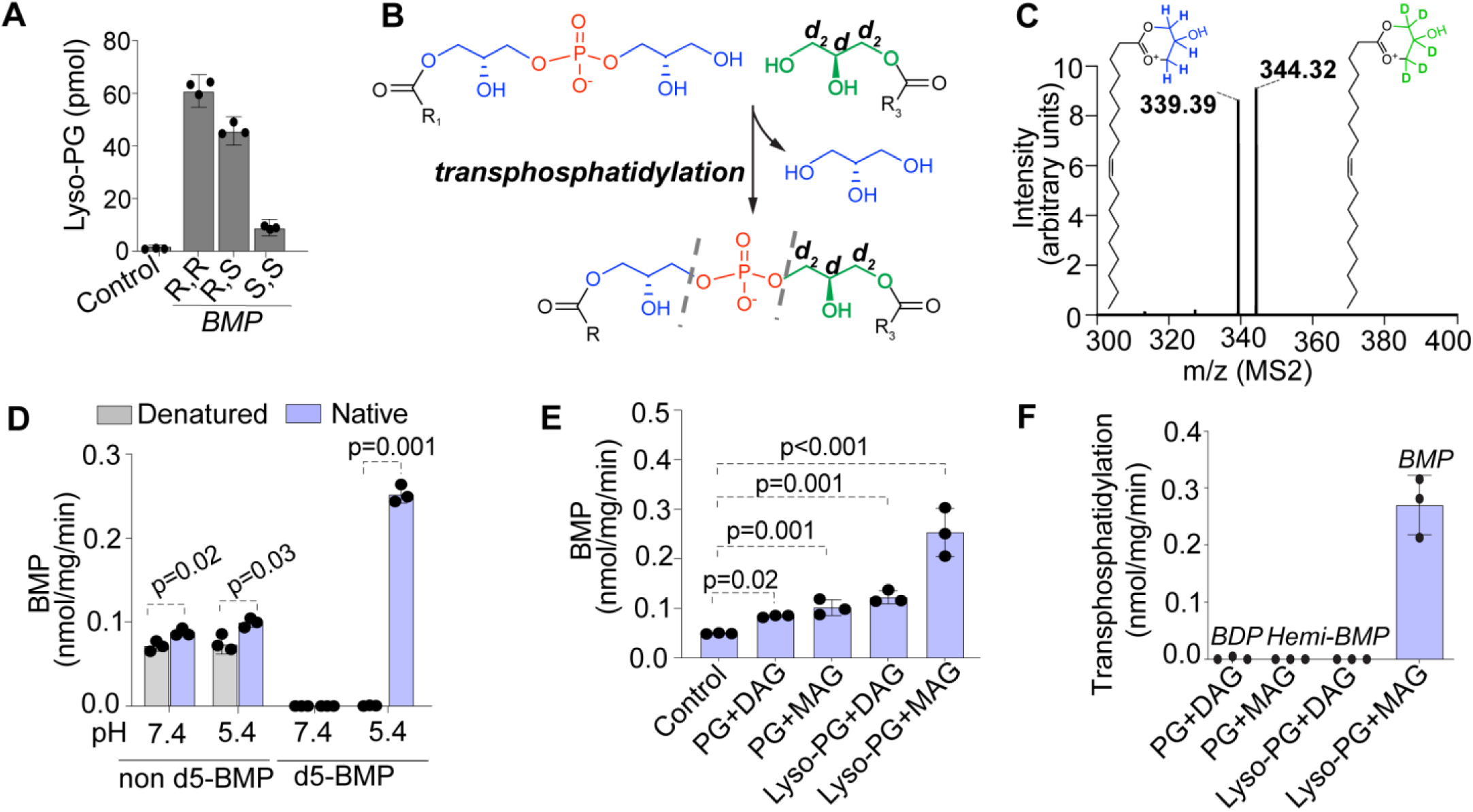
BMP is synthesized by transphosphatidylation of lyso-PG in mammalian lysosomes. (A) Stability of BMP stereoisomers incubated with lysosomal extracts of HMC3 cells. Data presented as mean ± S.D. (n=3 per group). Control, heat-denatured extract. BMP used in the reaction were: bis((*R*)-2-hydroxy-3-(oleoyloxy)propyl) phosphate as *R,R*-BMP, (*R*)-2-hydroxy-3-(oleoyloxy)propyl ((*S*)-2-hydroxy-3-(oleoyloxy)propyl) phosphate as *R,S*-BMP, and bis((*S*)-2-hydroxy-3-(oleoyloxy)propyl) phosphate as *S,S*-BMP. (B) Scheme of transphosphatidylation reaction where d5-containing MAG reacts with lyso-PG to generate d5-BMP. (C) MS2 fragment spectra depicting two fragments of d5-18:1/18:1 BMP; m/z=339.39 corresponds to an MAG-like fragment moiety without d5 glycerol, and m/z=344.32 corresponds to MAG moiety with d5 glycerol. (D) Relative transphosphatidylation (d5-BMP) and transacylation activities (non-d5-BMP) in lysosome extracts of HMC3 cells. pH 7.4 assays with DPBS buffer, and pH 5.4 with sodium-citrate buffer. (E) BMP synthesis activity in the lysosomes of HMC3 cells incubated with different lipid substrates. (F) Transphosphatidylation activities for different substrates in the lysosomes isolated from HMC3 cells. PG, R,rac-phosphatidylglycerol; DAG, diacylglycerol; lyso-PG, R,rac-lysophosphatidylglycerol; 1-MAG=1-(rac)monoacylglycerol. Data presented as mean ± S.D. (n=3 per group). *p*-values were calculated using Student’s t-test.

The synthesis of *S,S*-BMP from *R,S*-lyso-PG in mammals requires the stereochemical inversion of a chiral carbon in at least one glycerol, resulting in the *S*-stereo-conformation of both glycerol moieties. To test whether this stereo-inversion may involve a glycerol exchange or a re-arrangement of carbon substituents, we incubated lysosomal extracts with *R,rac*-lyso-PG and *sn*-1-(*rac*)-monoacylglycerol (MAG) containing a deuterium-labeled headgroup glycerol (18:1 d5-MAG) (**Figure 2B**). This resulted in the production of 18:1/18:1 d5-BMP (**Figure 2C,D**), which is the product of a transphosphatidylation reaction that exchanged one glycerol of lyso-PG with deuterated glycerol from MAG. The increase in d5-BMP synthesis was observed for incubations at pH 5.4 and not at neutral pH. In contrast, we found only minor increases in BMP synthesis at either pH via transacylation (generating BMP without deuterium label), where one acyl chain is transferred from a different glycerolipid to a lyso-PG molecule (**Figure 2D**). *In vitro* BMP synthesis activity was greatest for the substrates lyso-PG and MAG (**Figure 2E**). Other tested substrates did not yield products predicted for transphosphatidylation (**Figure 2F**). Thus, lysosomal extracts produce BMP by transphosphatidylation of lyso-PG and MAG substrates, which could lead to stereo-inversion of a glycerol moiety, a step required for *S,S*-BMP synthesis.

### PLD3 or PLD4 overexpression increase BMP levels in lysosomes

We sought to identify enzymes that catalyze the transphosphatidylation reaction to produce *S,S*-BMP. Previous studies showed that members of the phospholipase D (PLD) class of enzymes catalyze such reactions. For example, the bacterial phosphatidylserine synthase and cardiolipin synthases are members of the PLD enzyme family that use this catalytic mechanism (Centola et al., 2021; Tan et al., 2012a). We therefore focused on PLD enzymes as candidates. Mammals have six different PLD enzymes, and consistent with previous reports (Brown et al., 1998; Gonzalez et al., 2018a), three of them (PLD1, PLD3, and PLD4) localized to lysosomes when expressed in HMC3 cells (**Figure 3A,B**). In contrast, PLD2 localized to the cytoplasm, and PLD5 and PLD6 to mitochondria (**Figure S3A**). The catalytic activity of each of these enzymes is not completely understood, with for instance PLD3 reported as a phospholipase D (Nackenoff et al., 2021; Nibbeling et al., 2017) and an exonuclease (Gavin et al., 2021; Gavin et al., 2018).

**Figure 3.**
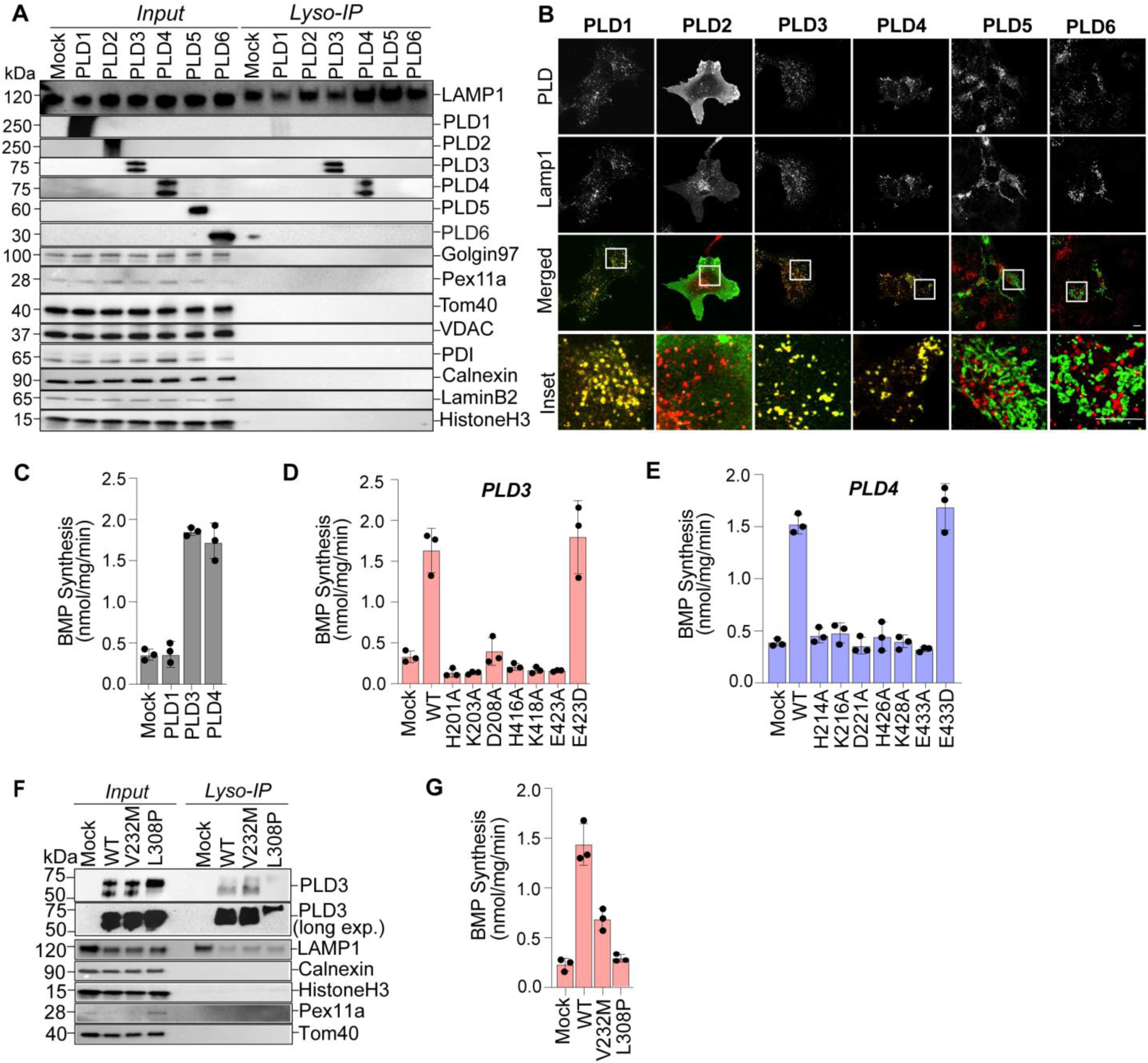
Expression of PLD3 or PLD4 in cells increased BMP levels. (A) Western blot of human PLD1-6 in lysosomes isolated after transfection of each PLD enzyme into HMC3 cells. Double bands in PLD3 and PLD4 blots correspond to full-length (∼75 kDa) and cleaved soluble form of the protein (∼60 kDa). (B) Confocal microscopy images of PLD1-6 expressed in HMC3 cells; PLD1-6 shown in green, LAMP1 shown in red. (C) BMP synthesis activities in lysosomes of PLD1-, PLD3-, or PLD4-transfected cells, compared with control cells. (D) BMP synthesis activities of PLD3 in lysosomes, compared with control cells and cells expressing mutants of the phosphodiesterase catalytic domain. (E) BMP synthesis activities of PLD4 in lysosomes, compared with control cells and cells expressing mutants of the phosphodiesterase catalytic domain. (F) Western blot of human patient mutants V232M and L308P PLD3 in lysosomes, compared with WT-PLD3. (G) Reduced BMP synthesis activities of V232M and L308P PLD3 in lysosomal lysates. Data are mean ± S.D. (n=3 per group).

To test whether lysosomal PLDs synthesize BMP, we overexpressed each of them in HEK293T cells expressing TMEM192-3xHA, which allows for isolation of lysosomes by rapid affinity purification (Abu-Remaileh et al., 2017). We tested the resultant lysosomal extracts for BMP synthesis activity and found that overexpression of PLD3 or PLD4, but not PLD1, increased BMP synthesis from lyso-PG and MAG substrates (**Figure 3C**).

PLDs have two conserved phosphodiesterase domains, each possessing a catalytic triad of histidine, lysine, and aspartic acid residues predicted to be essential for activity (**Figure S3B,C**). Mutations of any of these residues to alanine in PLD3 or PLD4 abolished lysosomal BMP synthesis activity (**Figure 3D,E**), but did not impair the expression or localization of the mutant proteins to lysosomes (**Figure S3D,E**). In contrast, a conservative change of E423D in PLD3 or E433D in PLD4 (aspartic acid is found at this position in other PLDs) had no detectable effect on expression, localization, or BMP synthesis activity (**Figure 3D,E and Figure S3D,E**).

In humans, mutations in PLD3 are implicated as causing dominant spinocerebellar ataxia 46 (Nibbeling et al., 2017; Sowmini et al., 2023). We tested one such missense mutation, PLD3 L308P, and found reduced lysosomal localization and essentially undetectable lysosomal BMP synthesis activity (**Figure 3F,G**). Immunoblotting also revealed decreased processing of the enzyme to its soluble form (**Figure 3F**). Another mutation, PLD3 V232M, increases the risk for Alzheimer’s disease (Yuan et al., 2022). PLD3 V232M localized normally to lysosomes but exhibited ∼50% reduced BMP synthesis activity (**Figure 3F,G**).

### Purified PLD3 or PLD4 catalyze *S,S*-BMP synthesis *in vitro* via a transphosphatidylation reaction

To test whether PLD3 or PLD4 are sufficient for catalyzing BMP synthesis, we overexpressed and purified each protein from HEK293FT cells (**Figure 4A,B**). Proteomic analyses of the purified protein samples revealed no contaminants likely to mediate BMP synthesis (including transphosphatidylation) reactions (**Table S1**). Either purified PLD3 or PLD4 catalyzed the synthesis of BMP from lyso-PG and MAG substrates *in vitro* (**Figure 4C,D**). In contrast, purified enzymes with mutations in the presumed catalytic sites (PLD3 H416A or PLD4 H214A) lacked BMP synthesis activity. Both PLD3 and PLD4 utilized *sn-*1*-*MAG or *sn-*2*-*MAG as substrates (**Figure 4C,D**). Additionally, PLD3 and PLD4 preferred *S,R*-lyso-PG (lyso-PG with acylation of the *S*-glycerol) compared with *R,racemic*-lyso-PG (lyso-PG with acylation of *R*-glycerol), although the enzymes could utilize either lyso-PG as a substrate. Neither purified enzyme was active with *R,racemic*-PG as a substrate.

**Figure 4.**
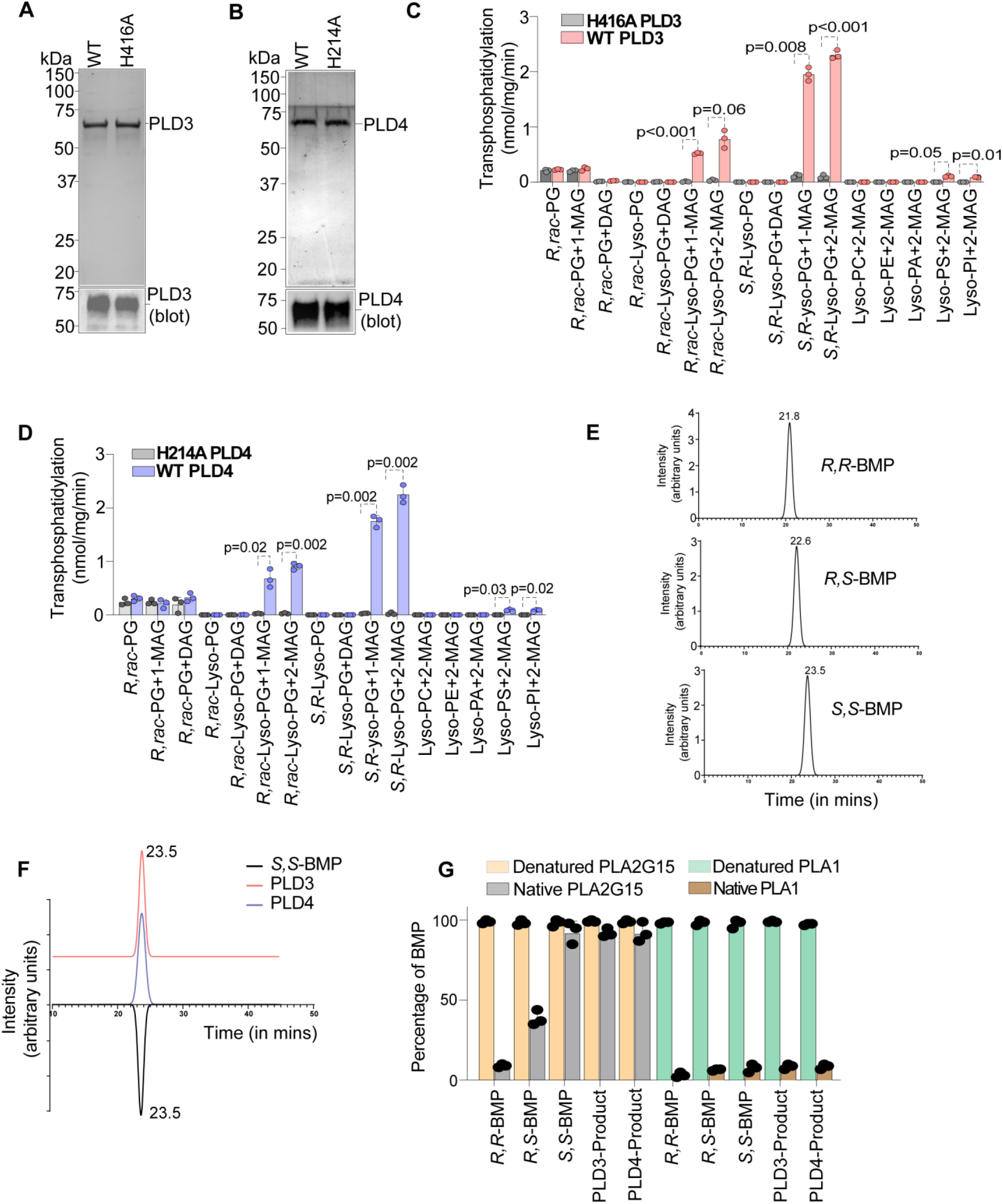
PLD3 or PLD4 was sufficient to catalyze *S,S*-BMP synthesis *in vitro*. (A) Coomassie-stained SDS-PAGE gel of purified recombinant WT and H416A-PLD3. (B) Coomassie-stained SDS-PAGE gel of purified recombinant WT and H214A-PLD4. (C) Transphosphatidylation activity of recombinant WT or H416A-PLD3 with various substrates. (D) Transphosphatidylation activity of recombinant WT or H214A-PLD4 with various substrates. R,rac-PG, (*R*)-2,3-bis(oleoyloxy)propyl (2,3-dihydroxypropyl) phosphate; 1-MAG, sn-1-(racemic) monoacylglycerol; 2-MAG, sn-2-(racemic) monoacylglycerol; DAG, sn-1,2-diacylglycerol; S,R-lyso-PG, (*S*)-2,3-dihydroxypropyl ((*R*)-2-hydroxy-3-(oleoyloxy)propyl) phosphate; R,rac-lyso-PG, 2,3-dihydroxypropyl ((*R*)-2-hydroxy-3-(acyloxy)propyl)phosphate; lysoPC, lysophosphatidylcholine; lysoPE, lysophophatidylethanolamine; lysoPA, lysophosphatidic acid; lysoPS, lysophosphatidylserine; and lysoPI, lysophosphatidylinositol. Data are mean ± S.D. (n=3 per group). *p*-values calculated by multiple t-test. (E) Elution trace of (R)-(-)-1-(1-naphthyl)ethyl isocyanate (R-NIC) derivatized BMP stereoisomers from a chiral column, demonstrating retention time differences of ∼1 min between *S,S-*, *R,S-* and *R,R*-BMP. (F) Elution profile of PLD3 and PLD4 products derivatized with R-NIC and S,S-BMP standard derivatized with R-NIC showing their co-elution profile. (G) Hydrolysis of standards of BMP stereoisomers and products of PLD3 and PLD4 when *S,R*-lyso-PG and 1-MAG were used as substrates by human PLA2G15 or *Aspergillus oryzae* PLA1 (Sigma-Aldrich #L3295). Denatured refers to heat-denatured protein and native refers to active enzyme used in the assay. Data presented as mean ± S.D. (n=3 per group).

To determine if the products of PLD3 and PLD4 reactions were the physiological *S,S*-BMP stereoisomer, we established methods to distinguish different BMP stereoisomers. First, we employed chiral derivatization [using (R)-(-)-1-(1-naphthyl)ethyl isocyanate (*R*-NIC)] and chromatographic separation to measure different BMP stereoisomers (**Figure 4E and Figure S3F**). Analyzing PLD reaction products, we found that derivatized products of PLD3 and PLD4 eluted at retention times (∼23.5 min) consistent with *S,S*-BMP rather than *R,R-*BMP (21.8 min) or *R,S*-BMP (22.6 min) (**Figure 4F**). Second, we assayed whether the products of PLD3 or PLD4 were substrates for PLA2G15, predicted to cleave glycerolipids in the *R*-configuration. BMP products of either PLD3 or PLD4 reactions were resistant to PLA2G15-mediated hydrolysis, whereas *R,R*-BMP or *R,S*-BMP were readily cleaved to lyso-PG. In contrast, incubation of the different BMPs with a control enzyme, PLA1 from *Aspergillus oryzae*, resulted in hydrolysis of all stereo-conformations of BMP, including the products of PLD3 and PLD4 (**Figure 4G and Figure S3G,H**). These results indicate that BMP synthesized by PLD3 or PLD4 is in the *S,S*-stereo-conformation (**Figure 4G and Figure S3G,H**).

### PLD3 and PLD4 deficiency in human cells resulted in decreased BMP levels, ganglioside accumulation, and other lysosomal abnormalities

We tested whether PLD3 and PLD4 are required for BMP synthesis in cells. HMC3 and HEK293T cells express PLD3, but not PLD4 (**Figure S4A**). To analyze PLD3/4 function in BMP synthesis, we therefore generated PLD3 knockout cells by CRISPR-Cas9-mediated genome engineering (**Figure 5A and Figure S4B**). BMP levels in PLD3-deficient HMC3 or HEK293T cells were 70–80% lower than those in control cells (**Figures 5B-D and Figure S4C,D**). BMP levels were restored by stably re-expressing PLD3 (“add-back” in **Figures 5B-D and Figure S4C,D**). Levels of lyso-PG or PG were similar in PLD3 knockout cells (**Figure S4E-F**).

**Figure 5.**
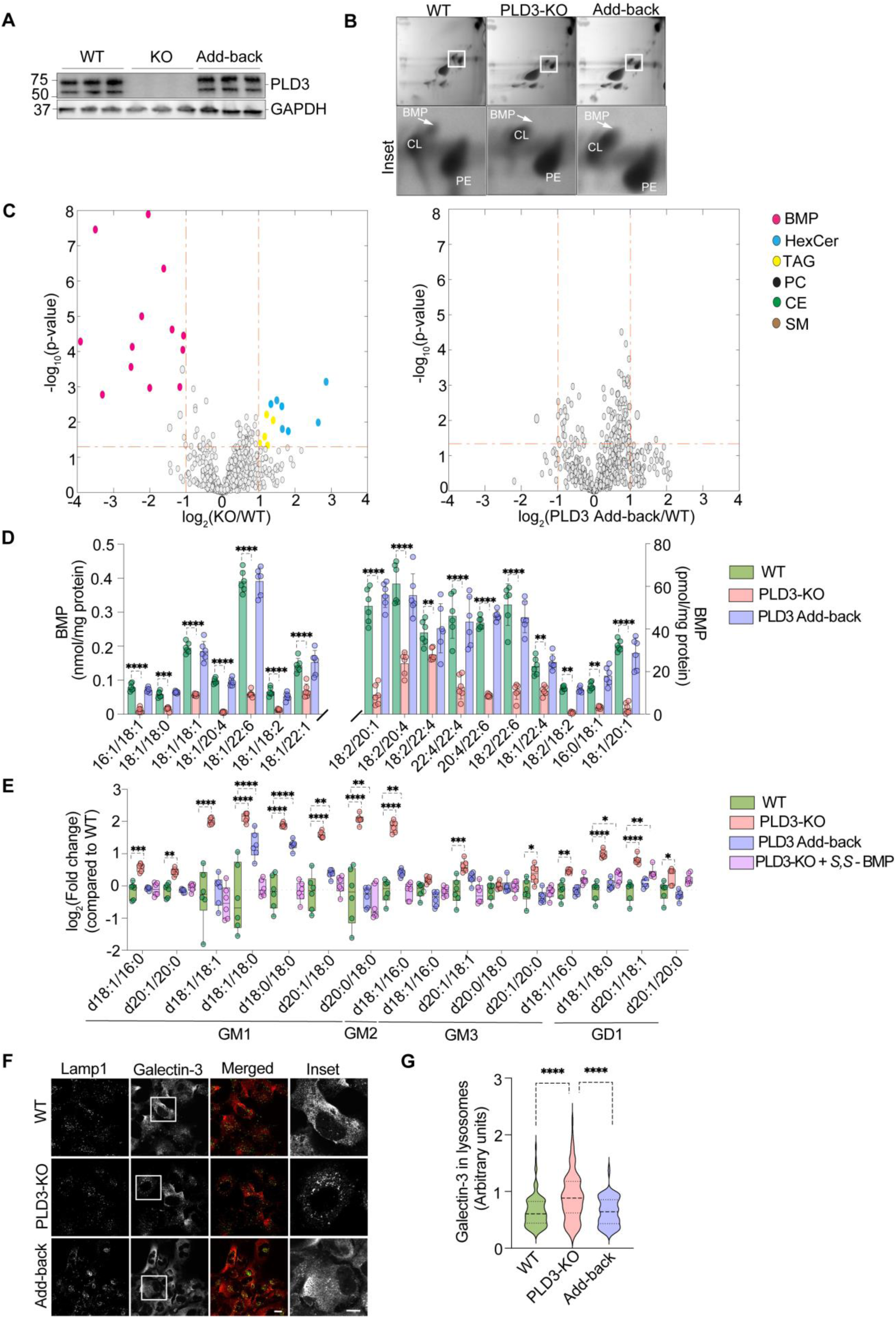
PLD3-knockout HMC3 cells exhibited reduced BMP levels, accumulation of gangliosides, and lysosomal abnormalities. (A) Western blot probing PLD3 presence in wildtype (WT), PLD3 knockout (KO), and add-back (i.e., PLD3 KO rescued by stable PLD3 expression) in HMC3 cells. (B) 2-D TLC demonstrating decreased BMP levels in PLD3 KO, compared with WT or add-back HMC3 cells. (C) Volcano plot representation of lipid measurements of PLD3 KO versus PLD3 WT and PLD3 add-back versus PLD3 WT HMC3 cells with log_2_-fold changes (ratio of relative abundance, x-axis) and log_10_ *p*-values (y-axis). BMP species were most decreased and are depicted as pink dots. Elevated lipids were mostly hexosylceramides (blue dots) and triacylglycerols (yellow dots). *p*-values calculated by two-sample t-test. (D) BMP species in WT, PLD3 KO and add-back HMC3 cells. (E) Ganglioside species in WT, PLD3 KO, add-back and PLD3 KO HMC3 cells fed with *S,S*-BMP. Data presented as mean ± S.D. (n=6 per group). \*\**p < 0.01*; \*\*\**p < 0.001*; \*\*\*\**p <0.0001* (two-way ANOVA multiple comparison with Dunnett’s correction). (F) Confocal microscopy images of cells stained with anti-galectin-3 (red) and anti-LAMP1 (green) antibodies, demonstrating increased galectin-3 localization in the lysosomes of PLD3 KO HMC3 cells. Scale bar =10 μm. (G) Quantification of galectin-3 signal (green) in the lysosomes (masked for red - LAMP1). \*\*\*\**p < 0.001*two-way ANOVA multiple comparison with Dunnett’s correction).

Since BMP is crucial for lipid degradation, reducing its levels may result in accumulation of lysosomal lipids, such as gangliosides (Abe and Shayman, 2009; Boland et al., 2022; Logan et al., 2021). Indeed, deletion of PLD3 from HMC3 or HEK293T cells resulted in accumulation of numerous species of gangliosides (**Figure 5E and Figure S4G**), and the ganglioside accumulation could be rescued either by adding back PLD3, or by feeding its reaction product *S,S*-BMP to cells (**Figure 5E and Figure S4G**). The transcript and protein levels of ganglioside metabolic enzymes were not significantly changed in PLD3 knockout cells or cells fed with BMP (**Figure S5A-C**). Additionally, *in vitro* activities of ganglioside catabolic enzymes were unchanged with PLD3 knockout (**Figure S5D**). These data indicate that ganglioside accumulation in PLD3 knockout cells is likely a consequence of low BMP levels, rather than due to an indirect effect, such as altered gene expression.

We also examined PLD3 knockout cells for lysosomal membrane defects. PLD3-knockout cells exhibited increased recruitment of galectin-3 to lysosomes, indicating a response to lysosomal damage that exposed glycans (**Figure 5F,G**). We also detected a modest reduction in lysosomal protease activity, assessed by fluorescent dequenching of DQ-BSA, in PLD3-knockout cells (**Figure S5E,F**). PLD3 has also been reported to function in nucleic acid degradation in lysosomes (Gavin et al., 2021; Gavin et al., 2018; Van Acker et al., 2023), and deficiency of PLD3 was shown to cause accumulation of mitochondrial DNA in lysosomes (Van Acker et al., 2023). At baseline, we found no differences in mitochondrial nucleic acid levels in the lysosomes of PLD3 knockout HMC3 cells, compared with wildtype (WT) cells **(Figure S5G)**, but induction of mitophagy by mitochondrial division inhibitor 1 (Mdivi-1) treatment increased mitochondrial DNA more in lysosomes of PLD3 knockout cells than in WT cells, as reported (Van Acker et al., 2023). However, for each of the genes assayed, this effect of PLD3 deficiency was reversed by *S,S*-BMP addition **(Figure S5G)**, consistent with this phenotype being a consequence of lowered BMP levels.

### PLD3 or PLD4 knockout mice exhibit reduced tissue levels of BMP and elevated ganglioside levels

*In vivo,* PLD3 and PLD4 appear to perform redundant functions because mice with either gene knocked out are viable but mice that lack both enzymes die before weaning (Gavin et al., 2021). Consistent with this hypothesis, systematic transcriptomic studies of mice (Wu et al., 2016) show that expression patterns of the enzymes vary greatly between tissues, with some tissues expressing predominantly one enzyme and other tissues having both enzymes.

According to systematic gene expression analyses in mice (Wu et al., 2016), PLD3 expression is highest in the central nervous system. To test the function of PLD3 in the brain, we studied global PLD3 knockout mice. Brains of PLD3 knockout mice had markedly reduced PLD3 mRNA levels and protein levels (**Figure 6A,B and S6A,B**). PLD3 knock out mice had a minor increase in levels of PLD4 levels but no change in levels of other detected PLDs (**Figure S6C**). In general, 8–12-week-old PLD3 knockout mice appeared healthy and exhibited normal behavior.

**Figure 6.**
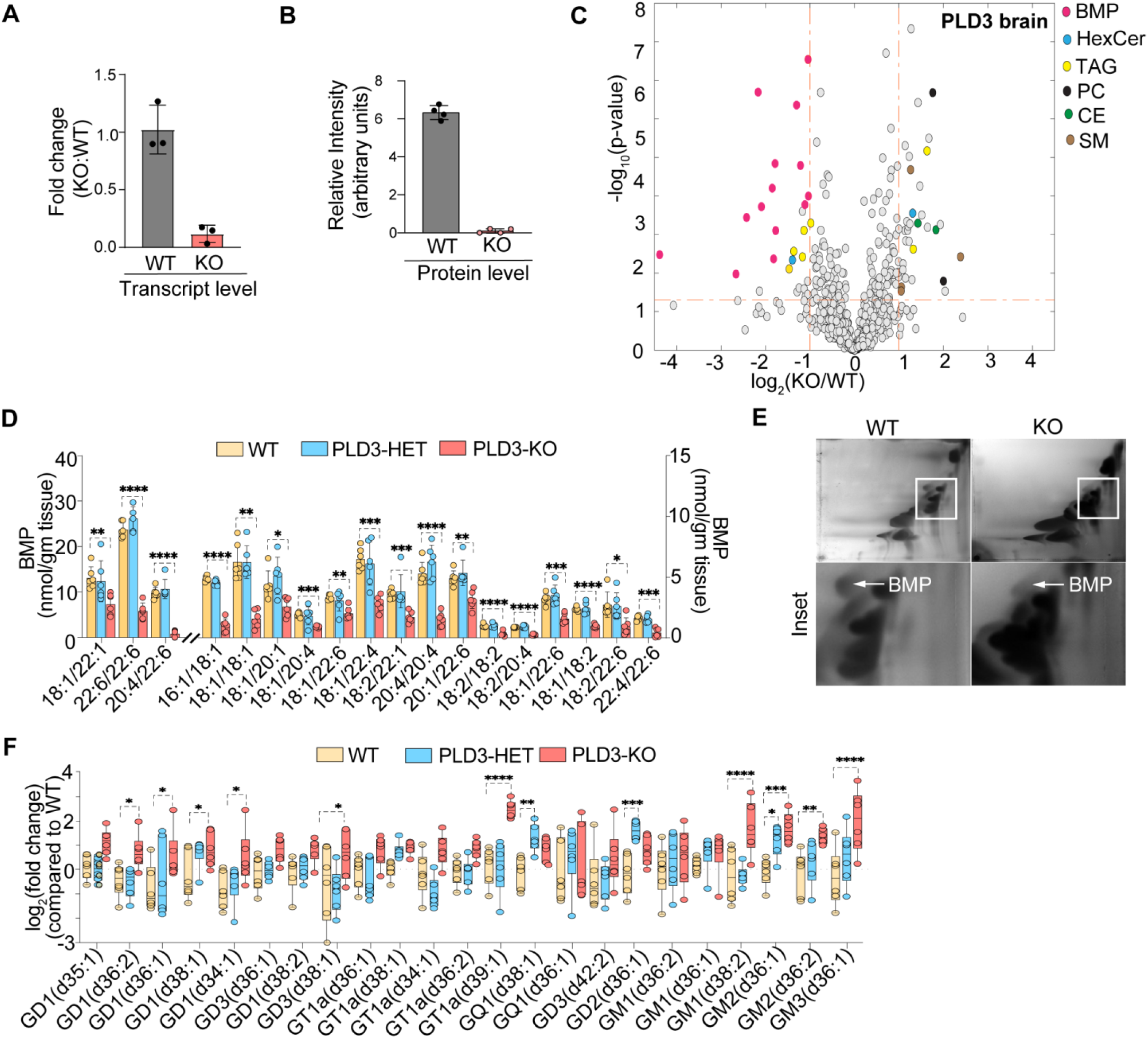
Reduced BMP and increased ganglioside levels in brains of PLD3 knockout mice. (A) RT-PCR data demonstrating loss of PLD3 expression in PLD3 KO mice brain. (B) Protein levels of PLD3 demonstrating loss of PLD3 protein expression in PLD3 KO mice brain. (C) Volcano plot representation of lipid measurements in the brains of PLD3 WT versus PLD3 KO mice with log_2_-fold changes (ratio of relative abundance, x-axis) and log_10_ *p*-values (y-axis). BMPs were most decreased depicted as pink dots. *p*-values calculated by two-sample t-test. (D) BMP species in brains of WT, PLD3 heterozygous (HET) and PLD3 KO mice. Data presented as mean ± S.D (n=6 per group). \**p < 0.05*; \*\**p < 0.01*; \*\*\**p < 0.001*; **** *p < 0.0001*. (two-way ANOVA multiple comparison with Dunnett’s correction). (E) 2-D TLC demonstrating a decrease in BMP levels in the brain of PLD3 KO mice, compared to their WT littermates. (F) Gangliosides species in the brains of WT, PLD3 HET and KO mice. Mice used were 3 males and 3 females of 2–3 months of age for each genotype. Data presented as mean ± S.D. (n=6 per group). \**p < 0.05*; \*\**p < 0.01*; \*\*\**p < 0.001*; \*\*\*\**p <0.0001* (two-way ANOVA multiple comparison with Dunnett’s correction).

Brains from 8–12-week-old PLD3 knockout mice had markedly reduced (∼70%) BMP levels and an accumulation of hexosylceramides in the brain, compared with wild-type or heterozygous littermate controls (**Figures 6C-E and Figure S6D**). In contrast, there were few changes of BMP lipid levels in peripheral tissues, such as the spleen, where PLD3 is less expressed (Wu et al., 2016) (**Figure S6E**). BMP deficiency results in defects in ganglioside degradation (Boland et al., 2022; Logan et al., 2021), and indeed brains of PLD3 knockout mice had accumulations of several different gangliosides, including GM1, GM2, GM3, GD1, and GD3 (**Figure 6F**).

In contrast to PLD3, PLD4 mRNA expression is highest in murine spleen and myeloid cells (Wu et al., 2016). To test the function of PLD4 *in vivo,* we studied PLD4 knockout mice generated by CRISPR-Cas9–mediated genome engineering. Knockout mice had markedly reduced PLD4 mRNA and protein levels in the spleen (**Figure 7A,B and S6F,G**). There were no differences in levels of other detected PLDs (**Figure S6H**).

**Figure 7.**
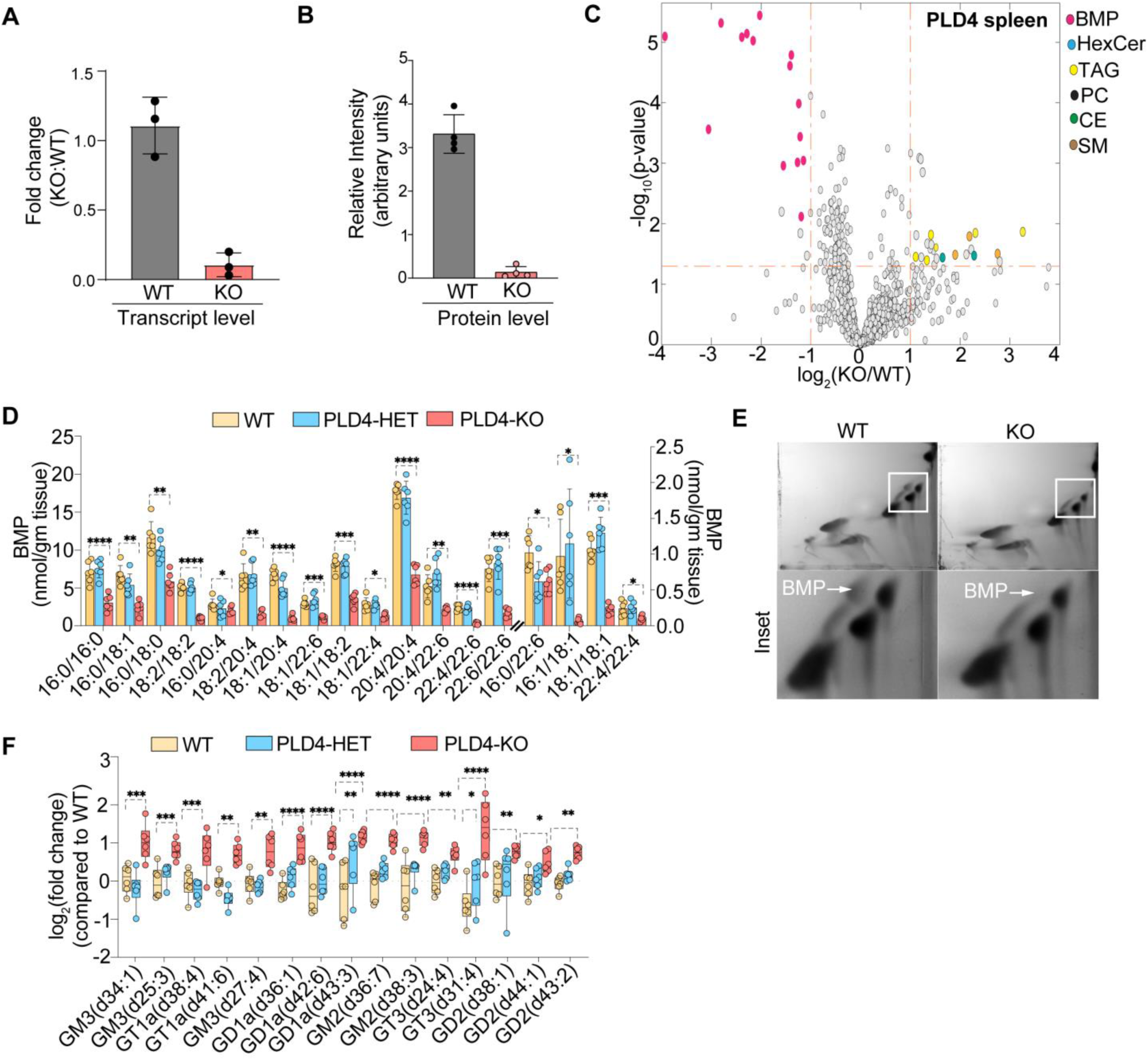
Reduced BMP and increased ganglioside levels in spleens of PLD4 knockout mice. (A) RT-PCR data demonstrating loss of PLD4 expression in PLD4 KO mice spleen. (B) Protein levels of PLD4 demonstrating loss of PLD4 protein expression in PLD4 KO mice spleen. (C) Volcano plot of lipid measurements in the spleen of PLD4 WT versus PLD4 KO mice with log_2_-fold-changes (ratio of relative abundance, x-axis) and log_10_ *p-*values (y-axis). BMPs were most decreased depicted as pink dots. *p-*Values calculated by two-sample t-test. (D) BMP species in the spleens of WT, PLD4 HET and PLD4 KO mice. Data presented as mean ± S.D. (n=6 per group). \**p < 0.05*; \*\**p < 0.01*; \*\*\**p < 0.001*; \*\*\*\**p <0.0001* (two-way ANOVA multiple comparison with Dunnett’s correction). (E) 2-D TLC demonstrating a decrease in BMP levels in the spleen of PLD4 KO mice, compared to WT littermates. (F) Gangliosides species in the spleens of WT, PLD4 HET and KO mice. Mice used were 3 males and 3 females of 2–3 months of age for each genotype. Data presented as mean ± S.D. (n=6 per group). \**p < 0.05*; \*\**p < 0.01*; \*\*\**p < 0.001*; \*\*\*\**p <0.0001* (two-way ANOVA multiple comparison with Dunnett’s correction).

Lipidomic analyses of PLD4 knockout mice revealed that PLD4 deficiency led to markedly decreased BMP levels (∼80%) in the spleen (**Figure 7C-E and Figure S6I**), whereas the deletion had little effect on BMP levels in the brain (**Figure S6J**). The reduction of BMP levels in the spleen correlated with a ∼twofold increase in the levels of GM3, GD1a, GD2, GT1, and GT3 gangliosides compared with WT or heterozygous litter-mate controls **(Figure 7F)**. In agreement with previous reports (Gavin et al., 2021; Gavin et al., 2018), PLD4 knockout mice exhibited splenomegaly (**Figure S6K,L**).

## DISCUSSION

*S,S*-BMP is the major species found in lysosomes and is crucial for lysosomal lipid degradation. How it is produced in the lysosome has been a long-standing mystery. Here we report that the lysosomal enzymes PLD3 and PLD4 catalyze *S,S*-BMP synthesis by mediating the crucial stereo-inversion step of the biosynthetic pathway. These enzymes were both sufficient and required to generate lysosomal BMP: purified enzymes were sufficient to catalyze *S,S*-BMP synthesis *in vitro* via a transphosphatidylation reaction, utilizing *S,R*-lyso-PG and MAG as substrates, and PLD3 and PLD4 were required for maintaining normal BMP levels in human cells and murine tissues.

Our findings provide a more complete picture of the *S,S-*BMP biosynthetic pathway. In the current model, the starting lipid is *R,S*-PG, a phospholipid predominantly synthesized in mitochondria (Kiyasu et al., 1963). How PG reaches the lysosome is unclear. In the lysosome, PG may be converted to lyso-PG by phospholipases, such as PLA2G15 (Chen et al., 2023), and lyso-PG could then serve as a precursor for BMP synthesis. In the final step, PLD3 or PLD4 perform the crucial stereo-inversion yield *S,S-*BMP, using *R*-lyso-PG and MAG as substrates (**Figure S7A**).

A recent report suggests that lysosomal CLN5 may be a BMP synthase, using two *R,racemic*-lyso-PG molecules as substrates in a transacylation reaction that generates *R,racemic*-BMP (Medoh et al., 2023). We purified CLN5 and found that *in vitro* it can use *R,racemic-*lyso-PG to synthesize *R,racemic-*BMP (**Figure S7B-C**), but not *S,S*-BMP. Possibly *R,S*-BMP or *R,R*-BMP made by CLN5 may be an intermediate that would be converted to lyso-PG by one of the abundant lysosomal phospholipases, such as PLA2G15 (**Figure S7A**). We question whether any transacylase is the crucial enzyme for BMP synthesis, since it would not mediate the glycerol stereo-inversion step required for *S,S*-BMP synthesis.

Previous studies reported that PLD3 and PLD4 provide exonuclease activities in the lysosome (Gavin et al., 2021; Gavin et al., 2018). While our studies do not exclude this possibility, we found no differences in nucleic acid levels in lysosomes of PLD3 knockout cells at baseline. As reported (Van Acker et al., 2023), we found that the induction of mitophagy increased mitochondrial DNA levels in PLD3 knockout lysosomes. However, this increase was reversed by addition of exogenous BMP, as was ganglioside accumulation (**Figure 5E**), suggesting that the loss of BMP is the primary defect in PLD3 deficiency. Alternatively, PLD3/4 enzymes may act as both lipases and nucleases, as reported for mitochondrial transmembrane PLD6 (Adachi et al., 2016; Baba et al., 2014; Choi et al., 2006; Huang et al., 2011; Ipsaro et al., 2012; Nishimasu et al., 2012; Zhang et al., 2016).

Our findings may be relevant to understanding neurodegenerative disease pathogenesis. The accumulation of gangliosides and other complex sphingolipids in lysosomes due to mutations of specific hydrolases has long been known to cause severe neurodevelopmental diseases, such as Gaucher’s, Tay-Sachs, and Sandhoff’s diseases (Hruska et al., 2008; Myerowitz and Costigan, 1988; Zampieri et al., 2009). More recently, accumulation of lysosomal lipids have been increasingly recognized as linked to adult neurodegenerative diseases, such as frontotemporal dementia (FTD) [*GRN* mutations (Boland et al., 2022)], motor neuron disease [*Vps54* mutations (Petit et al., 2020)], or Parkinson’s disease [*GBA* mutations (Hruska et al., 2008)]. The brain and neurons are particularly rich in complex sphingolipids, such as gangliosides (Puro et al., 1969), and degradation of these lipids is an important function of lysosomes in the brain (Breiden and Sandhoff, 2019; Sandhoff and Sandhoff, 2018). Thus, an understanding of the processes that maintain lysosomal BMP levels may illuminate a better understanding of disease mechanisms and provide novel avenues for their treatment. Consistent with this notion, reduced BMP levels in PLD3-deficient murine brain were associated with marked accumulation of gangliosides. This is similar to findings in progranulin-deficient FTD, where BMP deficiency leads to accumulation of gangliosides (Boland et al., 2022), and treatment with BMP rescued the ganglioside levels in cultured cells.

PLD3 mutations in humans have also been linked to human neurodegeneration. For example, the PLD3 V232M mutation is associated with increased Alzheimer’s disease risk (Heilmann et al., 2015; Yuan et al., 2022). We found that this mutation reduced BMP synthesis levels by more than 50%, suggesting that reduced levels of BMP may contribute to the increased Alzheimer’s disease risk. Another mutation, PLD3 L308P was reported to cause autosomal dominant spinocerebellar ataxia (SCA) 46 (Nibbeling et al., 2017). We found impaired PLD3 trafficking [consistent with a previous report (Gonzalez et al., 2018b)] and reduced BMP synthesis for the PLD3 L308P mutant protein. Thus, altered BMP synthesis may also contribute to some forms of SCA.

Our results for PLD3/4 enzymes playing an essential role for BMP synthesis and the elucidation of the BMP synthesis pathway in general should have broad implications, both for clarifying mechanisms of lysosomal lipid degradation and for deciphering the contribution of BMP and lysosomal lipid degradation to neurodegenerative diseases and other diseases of lysosomal storage.

## Acknowledgments

We thank members of Farese & Walther laboratory and Robert Zimmerman (U Graz) for helpful discussions and G. Howard for editorial assistance. Gifts of reagents are acknowledged in Materials and Methods. This work was supported by a grant from the Bluefield Project to Cure FTD (to R.V.F. and T.C.W.), a Human Frontiers Science Program fellowship grant (to S.S.), and postdoctoral fellowship grants from the Bluefield Project to Cure FTD (to S.S. and Y.A.). J.L.S. is supported by Medical Scientist Training Program grant to Weill Cornell/Rockefeller/Sloan Kettering Tri-Institutional MD-PhD program from the NIGMS-NIH (T32GM007739). T.C.W. is a Howard Hughes Medical Institute Investigator. We acknowledge NIH/NCI Cancer Center Support Grant (Core Grant P30 CA008748) to MSKCC. We thank Mara Monetti, Laura Tuffery, and Li Zhuoning (MSKCC-Proteomics Core) for discussions and assistance with proteomics, and Julian Dillon, Munoz Fabricio, and Kvin Lertpiriyapong (MSKCC-Center of Comparative Medicine and Pathology) for help with mouse husbandry.

## Author contributions

S.S., R.V.F., T.C.W. conceived the project, and R.V.F. and T.C.W. acquired project funding. S.S. and U.D. generated new reagents and performed biochemical experiments. J.L.S. helped with measurements of mRNAs and activities of ganglioside catabolic enzymes. S.S. performed microscopy and mass spectrometry experiments. Y.A. helped with ganglioside measurements. S.S., R.V.F., and T.C.W. co-wrote the manuscript with input from all authors.

## Materials and Methods

### Lead contact

Further requests and information concerning this study should be addressed to the lead contacts: Tobias C. Walther and Robert V. Farese Jr.

### Materials availability

All the reagents are available on request to lead contacts: Tobias C. Walther and Robert V. Farese Jr.

### Data and code availability

There was no new code written as the part of this study. Proteomics data are deposited in the ProteomeXchangeConsortium through PRIDE partner repository with dataset identifier PXD046505. All reagents, plasmids, antibodies, cell lines generated here are available from the Farese & Walther laboratory upon request. Other data generated are available from the corresponding authors on request. Source data are provided with this paper.

### Mice

PLD3 heterozygous (HET) on C57BL/6N and PLD4 heterozygous (HET) on C57BL/6J mice were obtained from the International Mouse Phenotyping Consortium (University of California, Davis). Heterozygous PLD3 or PLD4 mice were bred to generate homozygous PLD3 or PLD4 knockout (KO) and wildtype (WT) mice. Mice were genotyped for PLD3 alleles with the following primers: PLD3-forward 5′-TTCCTGTGTGCCTGTCTTGC-3′, PLD3-reverse 5′-AGCTCCTTTCTCCGTCCCTC-3′, and PLD3-transverse 5′-TCGTGGTATCGTTATGCGCC -3′. PCR amplification strategy yielded a 492-bp product for WT and a 178-bp product for KO allele. PLD4 mice were genotyped using the following primers: PLD4-forward 5′-CCAGCTTGAGGTAGCTTCTCATGG -3′, PLD4-reverse 5′-CCACTGATCAAGGCTGTTGCAG -3′, and PLD4-transverse 5′-GCTGACTAGGATAGGCTGGTTGGG -3′. PCR amplification yielded a 1162-bp product for WT and a 497-bp product for KO allele. All mice were housed in a pathogen-free barrier facility with a 12 h light/12 h dark cycle and *ad libitum* access to water and food. Animal procedures were approved by the Institutional Animal Care and Use Committee of the Memorial Sloan Kettering Cancer Centre on animals, following NIH guidelines.

### Cell culture

HMC3 (ATCC #CRL-3304), HEK293T (ATCC #CRL-3216) and their genetically manipulated versions were grown at 37°C, 5% (v/v) CO_2_ in Dulbecco’s Modified Eagles Medium (DMEM) (Invitrogen #11995-073), supplemented with 10% (w/v) heat-inactivated foetal bovine serum (FBS) (Gibco #10438026) and 1x penicillin-streptomycin (Gibco #15140163). HEK293FT cells (Invitrogen #R70007) were cultured and maintained in FreeStyle™ 293 Expression Medium (Gibco #12338026) supplemented with 5% (v/v) heat-inactivated FBS (Gibco #10438026) in a shaking incubator at 37°C, 5% (v/v) CO_2_. Cells were periodically tested for mycoplasma contamination with a PCR detection kit (abm #G238). All cell lines were verified by short tandem repeat profiling.

### RT-PCR

Total RNA was isolated from cells grown in six-well plates at 70–80% confluency with an RNA extraction kit (Qiagen #74104). cDNA was synthesized using a sensiFAST cDNA synthesis kit (Meridian Bioscience #BIO-65053). Genes were amplified for melting curves using Power SYBR^TM^ Green PCR Master Mix (Applied Biosystems^TM^, #4367659). Primers for the tested genes are listed in the reagents sheet.

### Molecular cloning

All plasmids used either were procured from GenScript or cDNA clones were obtained from Dharmacon – Mammalian Gene Collection (listed in the reagent sheet). All truncated or tagged versions of clones were generated using Q5 high-fidelity DNA polymerase (New England Biolabs #M0515) and NEB-Gibson Assembly (New England Biolabs #E2611S). All point mutations in PLD3 and PLD4 were generated using Q5-site directed mutagenesis kit (New England Biolabs #E0554S).

### Lentivirus production

HEK293T cells were transfected with pLJC5-hPLD3 or pLJC5-TMEM192-3xHA (Addgene #102930) envelope protein-pCMV-VSV-G (Addgene #8454), packaging plasmid - pCMV-dR8.2 dvpr (Addgene #8455) in a 6:4:1 ratio using PEI-max (Polysciences #24765-100). Virus-containing medium was harvested two days post-transfection, filtered through a 0.45-µm syringe filter, and stored at -80 ^°^C or used to transduce cells for stable cell line generation.

### Gene editing

CRISPR/Cas9 mediated gene editing of HEK293T and HMC3 cells was performed as described (Boland et al., 2022). Briefly, human PLD3 sgRNA (5′-3′) was cloned into pSpCa9(BB)-2A-Puro (PX459) V2.0 (Addgene #62988) as described (Ran et al., 2013). The sequences of sgRNA are as follows: forward – CTTGACGCCACGCTCGTAGG, reverse - CCTACGAGCGTGGCGTCAAG. Cells were transfected with 2 µg of plasmids with FuGene HD transfection reagent (Promega #E2312) overnight. Thereafter, cells were selected with puromycin (1 µg/mL) until all non-transfected control cells died (2–3 days). Selected cells were serially diluted to obtain knock out (KO) clones. To generate PLD3 rescue cell lines (also called PLD3 add-back cells), PLD3 KO HMC3 and HEK293T cells were transduced with lentivirus harbouring a PLD3 insert gene using polybrene at a final concentration of 10 µg/mL. TMEM192-3x-HA tagged (also called Lyso-Tagged) cells were generated as reported (Abu-Remaileh et al., 2017).

### Lipid incubations

Cells were plated in six-well dishes and grown to 70–80% confluency. Thereafter, medium was changed with fresh complete DMEM medium supplemented with 25 µM of phosphatidylglycerol, phosphatidylserine, phosphatidic acid, cardiolipin, or lysophosphatidylglycerol. Phosphatidylcholine and phosphatidylethanolamine were difficult to feed, and so, their concentrations were raised to 100 µM in the medium. Lipid-supplemented medium were prepared by drying lipids in a glass vial under N_2_ stream. Dried lipids were resuspended in complete DMEM using a water bath sonicator for ∼30 mins. Cells were maintained in lipid-supplemented medium for 6 h, and thereafter, cells were washed with ice-chilled Dulbecco’s phosphate-buffered saline (DPBS) (Gibco #14190144) six times and processed for lipid extraction or for immunofluorescence assays. For rescue experiments with BMP, knockout cells grown at ∼40% confluency were treated with 25 µM of *S,S*-BMP (prepared as described above) for 24 h, and thereafter, cells were washed with DPBS and processed for lipid extraction. All lipids used were procured from Avanti Polar Lipids.

### Immunostaining

Cells grown on glass cover slips were washed six times with DPBS, fixed with 4% (w/v) formaldehyde in PBS (BosterBio #AR1068) in DPBS for 15 min in the dark at room temperature (RT). Thereafter, cells were washed again three times with DPBS, permeabilized with permeabilization buffer containing 0.02% (w/v) saponin (Sigma-Aldrich #47036) and 1% (w/v) bovine serum albumin (Sigma-Aldrich #A6003) in DPBS at RT. Cells were incubated with blocking buffer containing 1% (w/v) bovine serum albumin in DPBS at RT, then incubated with primary antibody (1:100 dilution in blocking buffer) for 2 h at RT, washed three times with DPBS and incubated with secondary antibody (1:1000 dilution in blocking buffer) for 45 mins at RT. Cover slips were washed three times with DPBS and mounted on glass slides with Fluoromount Aqueous Mounting Medium (Sigma-Aldrich #F4680). Slides were dried at RT in the dark overnight before imaging. Information on the antibodies used is available in the reagent sheet.

### DQ-BSA dequenching assays

To measure lysosomal protease activity, DQ-BSA (Invitrogen #D12050) was dissolved at 1 mg/mL in DPBS by water-bath sonication. Cells were plated in glass-bottom 12-well plates and grown to 60– 70% confluency. Cells were washed with DPBS and fed with DQ-BSA (Invitrogen #D12050) at 100 µg/mL in complete DMEM medium for 4 h and treated with LysoTracker Red DND-99 (Invitrogen #L7528) for 15 min and subsequently imaged on a Nikon CSU-W1 SoRa microscope. Cells were maintained in a live cell chamber at 37°C, 85% humidity and 5% CO_2_.

### Microscopy and image analysis

Fluorescence images were acquired on a Nikon CSU-W1 SoRa spinning-disc microscope fitted with an ORCA-Fusion BT Digital CMOS camera (Hamamatsu #C15440-20IP). For cells, images were acquired through a 100x SR HP Apo 1.49 NA objective (Nikon #MRD71970). Green fluorophores (IgG-Alexa-488) were exited using a 488-nm laser and red fluorophores (IgG-Alexa-562) were exited using a 561-nm laser. Emission filters were for green signal = 525/50 and for red signal = 605/52. Images were processed and prepared for figures using Fiji-ImageJ. Signals in the lysosomes (e.g., BMP, DQ-BSA or galectin-3) were quantified by masking the signal for the lysosomal marker LAMP1 or LysoTracker Red DND-99 (Invitrogen #L7528).

### Western blot analysis

Cells were scraped into ice-chilled DPBS and pelleted at 1000 g for 2 min. Cell pellets were resuspended in DPBS, lysed by probe sonication (1 sec on, 3 sec off, 60% amplitude), boiled with 4X loading dye at 95°C for 30 min for denaturation. Denatured proteins were separated on 4–15% gradient SDS-PAGE gels at 120 V and transferred to methanol-activated PVDF blotting membrane at 60 V for 3 h at 4°C. Thereafter, PVDF membranes were blocked with 5% (w/v) skim milk with 0.1% (v/v) Tween-20 in DPBS (blocking buffer) for 1 h at RT. Blocked membranes were incubated with primary antibodies (1:1000 diluted in blocking buffer) overnight at 4° C, washed three times with 0.1% (w/v) Tween-20 in DPBS (washing buffer), incubated with secondary antibody (Santa Cruz Biotechnology #sc516102 #sc2357 - 1:5000 diluted in blocking buffer) for 1 h at RT. Membranes were washed three times with washing buffer and developed with ECL chemiluminescence substrates and imaged on a BioRad ChemiDoc Imager.

### Lysosome purifications

Lysosomes were immunoisolated as reported (Abu-Remaileh et al., 2017). TMEM192-3X HA-tagged cells from one 150-mm plate at 70% confluency were scraped into ice-chilled DPBS supplemented and pelleted at 1000 g for 2 min. The cell pellet was Dounce-homogenized in 1 mL ice-chilled KPBS buffer containing 136 mM KCl, 10 mM KH_2_PO_4_ adjusted to pH 7.25 with KOH. Cell lysates were spun down at 1000 g for 2 min. Supernatant was collected and incubated with anti-HA magnetic beads, washed and pre-equilibrated with KPBS buffer for 15 min at 4° C. Beads were washed with excess KPBS buffer 10 times. Proteins were eluted from the beads using lysis buffer containing 50 mM Tris, 150 mM NaCl, 1 mM EDTA, 1% (v/v) Triton X-100 adjusted to pH 7.4.

### Recombinant protein expression and purification

HMC3 or HEK293T cells were grown at ∼50% confluency in complete DMEM medium and transfected with pSPORT6 expression plasmids carrying human versions of PLD1, PLD2, PLD3, PLD4, PLD5, and PLD6 genes, using FuGene HD transfection reagent (Promega #E2312) for 24 h and thereafter processed for immunostaining or for lysosomal isolations and western blots. For PLD3, PLD4, PLA2G15 and CLN5 purifications, HEK293FT cells were grown at 1 x 10^6^ cells/mL and transfected with C-terminal FLAG-tagged WT or active-site mutants cloned in pcDNA3.1 using PEI-max (Polysciences #24765-100). Cells were treated with 10 mM sodium butyrate for ∼16 h after transfection and cultured for two additional days. Cells were collected by centrifugation at 2500 g for 15 min, and cell pellets were rinsed with DPBS. Cell pellets were resuspended in 50 mM Tris (pH 7.5), 200 mM NaCL, 5 mM MgCL_2_, 1 mM EDTA, 5 mM TCEP (resuspension buffer) and lysed by probe sonication (60% amplitude, 1 sec on, 3 sec off for 10 min). Lysates were centrifuged at 25,000 g for 1 h at 4° C. Supernatants were collected, syringe filtered with a 0.45-µM syringe filter, and incubated with anti-FLAG beads (Sigma-Aldrich #A2220) pre-equilibrated with resuspension buffer without TCEP. Beads were washed 10 times with 10 column volumes of resuspension buffer without TCEP. Proteins were eluted with 50 µg/mL of 1x FLAG peptide (Sigma-Aldrich #F3290), concentrated using 30 kDa centricons (Millipore #UFC9030). Concentrated proteins were buffer exchanged with sodium citrate buffer, pH 5.4, using button dialysis, followed by centrifugation at 50,000 g for 30 min to pellet any aggregates. Protein concentration in the supernatant fraction was accessed by nanodrop spectrophotometry, and purity of the protein was accessed using 10% Coomassie-stained SDS-PAGE gel and label-free mass spectrometry.

### *In vitro* enzymatic assays

For BMP synthesis assays, lipid substrates were prepared by resuspending the required amount of lipids in sodium citrate buffer at pH 5.4 using water-bath sonication to a final concentration of 50 μM of phospholipids and 25 μM of mono- or diacylglycerol (Avanti Polar Lipids). Substrates were incubated with extracts of affinity isolated lysosomes on anti-HA magnetic beads (3–5 μg of lysosomal proteins) or 1 μg of purified enzyme (final volume = 100 μL) for 45 min, 37 °C with constant shaking in glass vials. Thereafter, the reaction was quenched with 250 μL of 2:1 chloroform:methanol spiked with 14:0/14:0 BMP as internal standard. The mixture was vortexed and spun at 1000 g for 5 min, and the bottom organic layer was collected in a separate glass vial. Thereafter, the aqueous fraction was acidified with 2 μL of formic acid, vortexed and extracted again with 250 μL of chloroform. Organic layers from both steps were pooled, dried under a N_2_-stream and analyzed by LC-MS/MS. BMP formation was monitored and normalized to internal standards, the amount of protein used and the time of the reaction.

To test the stability of BMP stereoisomers in lysosomes 100 μM of 18:1/18:1 BMP stereoisomers were incubated with 5 μg of lysosomal lysates for 1 h at 37 °C with constant shaking in glass vials. To test the stability of BMP synthesized by PLD3 or PLD4, enzymatic products were incubated with 1 μg of PLA2G15 or 5 μg of phospholipase A1 from *Aspergillus oryzae* (Sigma-Aldrich #L3295) for 15 mins at 37 °C. This incubation time was reduced to accommodate for the low amount of BMP synthesized by PLD3 or PLD4 compared to their commercially available synthetic standards. Reaction was quenched with 250 μL of 2:1 chloroform:methanol; and lipids were extracted as described for the BMP synthesis assay. Hydrolysis of BMP stereoisomer standards was monitored by TLC and reaction rates were measured by LC-MS/MS. PLA2G15 used in the reactions was obtained by overexpressing C-terminally FLAG-tagged human PLA2G15 in HEK293FT cells and affinity-isolation using anti-FLAG magnetic beads (Sigma-Aldrich #M8823).

To test for lysosomal catabolic activity (e.g., ganglioside catabolism), a fluorescence plate reader assay was setup as described (Boland et al., 2022). Artificial substrate mimetics for lysosomal enzymes were dissolved in DMSO to stock concentrations of 100 μM. Cells were Dounce-homogenized in ice-cold DPBS and probe sonicated for 1 min (60% amplitude, 1 sec on, 3 sec off). Protein concentrations were estimated using Pierce BCA Protein Assay Kit (Pierce #23225) with BSA as control for standard curves. Lysates (25 μg) were incubated with 1 mM of fluorogenic substrates in sodium citrate buffer (total volume = 100 μL) at pH 5.4 at 37 °C for 30 min. Thereafter, reactions were quenched with 300 μL glycine-buffer, pH 10.4. Fluorescence was measured in a Tecan Spark multimode microplate reader with excitation and emission set at 350 and 440 nm. A standard curve was drawn using 4-methylumbelliferone (Sigma-Aldrich #M1381) as standards dissolved in reaction buffer + quenching buffer in ratios used for the assay. Substrates used were as follows: 4-methylumbelliferyl-β-D-galactoside for β-galactosidase, 4-methylumbelliferyl-β-D-glucopyranoside for β-glucocerebrosidase, 4-nethylumbelliferyl-2-acetamido-2-deoxy-β-D-glucopyranoside for β-hexosaminidase, 7-(α-D-glucopyranosyloxy)-4-methyl-2H-1-benzopyran-2-one for α-glucosidase, 7-(α-D-galactopyranosyloxy)-4-methyl-2H-1-benzopyran-2-one for α-galactosidase, 7-[[2-(acetylamino)-2-deoxy-α-D-galactopyranosyl]oxy]-4-methyl-2H-1-benzopyran-2-one for β-galactopyranosaminidase activity, hexadecanoic acid, 4-methyl-2-oxo-2H-1-benzopyran-7-yl ester for lysosomal acid lipase, and 4-methyl-2-oxo-2H-1-benzopyran-7-yl 5-(acetylamino)-3,5-dideoxy-α-neuraminic acid for neuraminidases. All artificial substrates were procured from Cayman Chemicals.

### Preparation of *S,R*-lyso-PG

*R,S*-BMP (100 μM) was incubated with 5 μg of human PLA2G15 at 37 °C in glass vials with constant shaking. The reaction was monitored by running small aliquots on TLC. The reaction was quenched and extracted with excess 2:1 (v/v) chloroform:methanol. Extracts were dried under an N_2_ stream, and the lyso-PG mixture was separated on a C18 reverse phase column (Pheneomenex, 5 µm, 50 x 4.6 mm) using the following solvents: Solvent A= 95:5 (v/v) H_2_O/MeOH + 0.1% (v/v) NH_4_OH, solvent B= 60:35:5 (v/v) IPA/MeOH/H_2_O + 0.1% (v/v) NH_4_OH. The liquid chromatography (LC) gradient was 30 mins, with a stable flow rate of 0.5 mL/min, starting 5% B for 5 min, linear gradient of B from 5– 100% over 23 min, 100% B from min 23–25, and re-equilibration with 5% B for 5 min. The isolated lyso-PG was pooled and dried under N_2_, weighed and used for BMP synthesis assays.

### Chiral derivatization and separation of BMP stereoisomers

Standards of BMP stereoisomers or enzymatic products were mixed with derivatization agent (R)-(-)-1-(1-naphthyl)ethyl isocyanate (Sigma-Aldrich #220442) in a ratio of 1:10 in 1 mL of dichloromethanol (VWR BDH23373.400) and 50 μL of pyridine (Sigma-Aldrich #270970) overnight (14–16 h) at RT. The reaction mixture was dried under a stream of N_2_ and extracted with 2:1:1 chloroform:methanol:water. The bottom organic layer was collected and separated on YMC Chiral ART Cellulose-SZ, analytical column (YMC America #KSZ99S03-2502WT). Solvent A = 60:40 (v/v) ACN/water + 0.1% (v/v) formic acid + 10 mM ammonium formate, Solvent B = 90:10 (v/v) IPA: ACN + 0.1% (v/v) formic acid + 10 mM ammonium formate. The LC gradient was 45 min, with a stable flow rate of 0.2 mL/min, starting with 40% B for 5 min, linear gradient of 40–70% B for min 5–30, maintained at 100% B for mins 30–33, re-equilibrated with 40% B for min 33–45 at 0.3 mL/min. The column temperature was maintained at 10 °C. MS parameters were as described in LC-MS/MS analysis of lipids section.

### Lipid extraction

Lipids were extracted from cells or tissues using established protocols (Kelkar et al., 2019). Briefly, cells were lysed in DPBS using a probe sonicator, or tissues were Dounce-homogenized in DPBS, and lipids were extracted with a chloroform and methanol mixture, with spiked-in lipid splash mix and/or BMP 14:0/14:0, as internal standards. The final ratio of chloroform:methanol:DPBS in the extraction mixture was 2:1:1. The extraction mixture was vortexed and spun down at 1000 g for 10 min. The bottom organic phase was collected in a glass tube. The top aqueous phase was acidified with 0.5% (v/v) formic acid, vortexed and re-extracted with chloroform by spinning down as in the previous step. Both organic phases were pooled and dried under a stream of N_2_ and analysed by LC-MS/MS.

### Ganglioside extraction

Tissues or cells were Dounce-homogenized with ice-chilled DPBS, and methanol was added to the final ratio of 90:10 methanol:water. The extraction mixture was spun down at 1000 g for 20 min to pellet proteins, and the solvent layers containing gangliosides and other metabolites were collected in glass tubes and dried under an N_2_ stream. Dried extract was reconstituted in 1 mL of LC-MS-grade water and desalted by Sola HRP SPE 30mg/2MI 96-well plate 1EA (Thermo Scientific #60509-001). Desalting cartridges were cleaned three times with 1 mL of methanol, equilibrated three times with LC-MS-grade water, and then the extracts dissolved in water were loaded onto the cartridge and washed three times with water, and finally, gangliosides were eluted by 3 mL of methanol. Eluates were dried under N_2_ and reconstituted in 300:150:50 of MeOH:water:chloroform.

## TLC for lipids

Extracted lipids were separated on high-performance thin layer chromatography (Sigma-Aldrich #1056310001) in the first dimension using 70:30:3:2 (v/v/v/v) chloroform:methanol:water:concentrated ammonia as the running phase. For BMP separation, plates were dried at RT for 1 hour and run in the second dimension using 65:35:5 (v/v/v) of chloroform:methanol:water as the running phase. Plates were dried and sprayed with 5% (w/v) copper sulfate dissolved in 25% (v/v) ethanol and 8.5% (v/v) phosphoric acid, followed by charring at 60 °C for 1 h. Plates were imaged on BioRad ChemiDoc Imager.

### LC-MS/MS analysis of lipids

All samples were analysed on a Vanquish UHPLC (Thermo Scientific) coupled to a Orbitrap Exploris 240 mass spectrometer (Thermo Scientific #BRE725535). For routine lipid analysis, extracts were separated on a C18 reverse phase column (Pheneomenex, 5 µm, 50 x 4.6 mm #00B-4435-E0) as described (Kelkar et al., 2019). Briefly, for positive mode: solvent A= 95:5 (v/v) H_2_O/MeOH + 0.1% formic acid + 10 mM ammonium formate, solvent B= 60:35:5 (v/v/v) isopropanol (IPA)/MeOH/H_2_O + 0.1% (v/v) formic acid + 10 mM ammonium formate. For negative mode, solvent A= 95:5 (v/v) H_2_O/MeOH + 0.1% (v/v) NH_4_OH, solvent B= 60:35:5 (v/v) IPA/MeOH/H_2_O + 0.1% (v/v) NH_4_OH. LC gradient was 60 min, with stable flow rate of 0.5 mL/min starting 5% B for 5 min, linear gradient of B from 5–100% over 50 min, 100% B from min 50–55, and re-equilibration with 5% B for 5 min. The column temperature was maintained at 55 °C. For BMP and PG, separation was performed on a C-30 reverse phase column (Thermo Scientific #078664, 3 µm 250 x 2.1 mm). Solvent = 60:40 (v/v) ACN/water + 0.1% (v/v) formic acid + 10 mM ammonium formate, Solvent B = 90:10 (v/v) IPA: ACN + 0.1% (v/v) formic acid + 10 mM ammonium formate. The LC gradient was 45 min, with a stable flow rate of 0.2 mL/min starting 40% B, linear gradient of 45–55% over mins 1–7, maintained at 65% B for mins 8–12, linear gradient of 65–70% B for mins 12–30, maintained at 100% B for mins 30–33, re-equilibrated with 40% for mins 33–45. Column temperature was maintained at 30 °C. MS parameters were as follows: - heated ESI mode of ionization, sheath gas = 40 units, aux gas = 8 units, sweep gas =1 units, spray voltage = static, positive ion =3,400 V, negative ion = 2500 V, ion transfer tube temp= 320 °C, vaporizer temp=275 °C. Orbitrap resolution at MS1 was set to 120000, scan range 250–1800, RF lens = 60%, AGC target = standard, Orbitrap resolution at MS2 was set to 30000. EASY-IC^TM^ was used for internal calibration. For general lipidomics, lipids were searched and aligned for annotation in LipidSearch 5.0 (Thermo Scientific #OPTON-30880) with precursor mass error tolerance of 5 ppm and product mass error tolerance of 10 ppm, retention time deviation tolerance of 0.5 min. For PG and BMP analysis, the orbitrap was set up with an unscheduled targeted inclusion mass list corresponding to PG and BMP. PG and BMP peaks were identified using Excalibur 3.1 (Thermo Scientific #OPTON-30965), the area under the curve was normalized to the area under the curve for respective internal standards and amount of protein/wet weight of tissue used.

### LC-MS/MS analysis of gangliosides

For LC-MS/MS analysis of gangliosides, samples were analyzed on a Vanquish UHPLC (Thermo Scientific) coupled to a Orbitrap Exploris 240 mass spectrometer (Thermo Scientific #BRE725535). Extracts were separated on a Kinetex HILIC column (Phenomenex #00D-4461-AN, 2.6 µm,100 x 2.1 mm). Solvent A= acetonitrile with 0.2% (v/v) acetic acid, solvent B = water with 10 mM ammonium acetate, pH 6.1, adjusted with acetic acid. The column temperature was maintained at 50 °C. The LC gradient started with a constant flow rate of 0.6 mL/min with 12.3% B at 0 min, linear gradient of 12.3% B to 22.1% B for min 1–15. The column was equilibrated between runs by 12.3% B for 5 mins. The spray voltage was set to −4.5 kV, and the heated capillary and the HESI were held at 300 °C and 250 °C, respectively. MS parameters were as follows: - heated ESI mode of ionization, sheath gas = 40 units, aux gas = 5 units, sweep gas =1 units, spray voltage = static, negative ion =-4,500 V, negative ion = 2500 V, ion transfer tube temp= 300 °C, vaporizer temp=250 °C. Orbitrap resolution at MS1 was set to 120000, scan range 250–1800, RF lens = 60%, AGC target = standard, Orbitrap resolution at MS2 was set to 30000. EASY-IC^TM^ was used for internal calibration. Analysis was performed in Excalibur (Thermo Scientific #OPTON-30965). The area under the curve was normalized to GM3-d5 standards, followed by normalization to amount of protein/wet weight of tissue used.

### Proteomics

Cellular lysates (50 μL) at 1 mg/mL were mixed with 50 μL of 8 M urea, 50 mM EPPS. The sample was then incubated with 5 mM TCEP for 30 min at 25 °C at 1000 rpm. Then it was incubated with 10 mM IAA during 30 min in the dark at RT before adding 5 mM DTT for 15 min at RT. The sample was digested with Lys-C with a 1:200 ratio enzyme:protein for 1 h at 25 °C at 1000 rpm. The sample was finally diluted half with 50 mM ammonium bicarbonate before digesting with trypsin with a 1:100 ratio protein:enzyme at 37 °C overnight. After the overnight incubation, the samples were desalted using C18 stage tips. Peptides were separated on a C18 reverse phase column (25-cm length, 75-μm diameter and 1.7-μm particle size from IonOptic Aurora 3 #1801220) using a gradient of 2–95% B over 90 min at 50 °C with a flow rate of 300 nL/min using a NanoElute 2 system (Bruker). Solvent A= 0.1% (w/v) formic acid in HPLC-grade water; solvent B = 99.5% (v/v) acetonitrile + 0.1% (v/v) formic acid. MS data were acquired on a timsToF (Bruker) with a captive spray source in data-independent acquisition mode. The mass range was set from 100 to 1700 m/z, the ion mobility range was set from 0.60 V.s/cm2 (collision energy 20eV) to 1.6 V.s/cm2 (collision energy 59 eV) with a ramp time of 100 ms and an accumulation time of 100 ms. Dia-PASEF settings were mass range 400.0 Da to 1201.0 Da, mobility range 0.60–1.60 and a cycle time estimate of 1.80 sec. The dia-PASEF windows were set with a mass width of 26.00 Da, mass overlap 1.00 Da, mass steps per cycle 32. Dia windows are as listed:

| #MS Type | Cycle Id | Start IM [1/K0] | End IM [1/K0] | Start Mass [m/z] | End Mass [m/z] | CE [eV] |
| --- | --- | --- | --- | --- | --- | --- |
| MS1 | 0 | - | - | - | - | - |
| PASEF | 1 | 0.91 | 1.6 | 800 | 826 | - |
| PASEF | 1 | 0.6 | 0.904 | 400 | 426 | - |
| PASEF | 2 | 0.9236 | 1.6 | 825 | 851 | - |
| PASEF | 2 | 0.6 | 0.9174 | 425 | 451 | - |
| PASEF | 3 | 0.9372 | 1.6 | 850 | 876 | - |
| PASEF | 3 | 0.6 | 0.9308 | 450 | 476 | - |
| PASEF | 4 | 0.9507 | 1.6 | 875 | 901 | - |
| PASEF | 4 | 0.6 | 0.9442 | 475 | 501 | - |
| PASEF | 5 | 0.9643 | 1.6 | 900 | 926 | - |
| PASEF | 5 | 0.6 | 0.9576 | 500 | 526 | - |
| PASEF | 6 | 0.9779 | 1.6 | 925 | 951 | - |
| PASEF | 6 | 0.6 | 0.971 | 525 | 551 | - |
| PASEF | 7 | 0.9915 | 1.6 | 950 | 976 | - |
| PASEF | 7 | 0.6 | 0.9844 | 550 | 576 | - |
| PASEF | 8 | 1.0051 | 1.6 | 975 | 1001 | - |
| PASEF | 8 | 0.6 | 0.9979 | 575 | 601 | - |
| PASEF | 9 | 1.0186 | 1.6 | 1000 | 1026 | - |
| PASEF | 9 | 0.6 | 1.0113 | 600 | 626 | - |
| PASEF | 10 | 1.0322 | 1.6 | 1025 | 1051 | - |
| PASEF | 10 | 0.6 | 1.0247 | 625 | 651 | - |
| PASEF | 11 | 1.0458 | 1.6 | 1050 | 1076 | - |
| PASEF | 11 | 0.6 | 1.0381 | 650 | 676 | - |
| PASEF | 12 | 1.0594 | 1.6 | 1075 | 1101 | - |
| PASEF | 12 | 0.6 | 1.0515 | 675 | 701 | - |
| PASEF | 13 | 1.073 | 1.6 | 1100 | 1126 | - |
| PASEF | 13 | 0.6 | 1.0649 | 700 | 726 | - |
| PASEF | 14 | 1.0865 | 1.6 | 1125 | 1151 | - |
| PASEF | 14 | 0.6 | 1.0783 | 725 | 751 | - |
| PASEF | 15 | 1.1001 | 1.6 | 1150 | 1176 | - |
| PASEF | 15 | 0.6 | 1.0918 | 750 | 776 | - |
| PASEF | 16 | 1.1137 | 1.6 | 1175 | 1201 | - |
| PASEF | 16 | 0.6 | 1.1052 | 775 | 801 | - |

### Mitophagy induction and nucleic acid measurement in lysosomes

Mitochondrial DNA in the lysosomes was quantified as described previously (Van Acker et al., 2023). Briefly, lyso-tagged WT and PLD3 knock out HMC3 cells were treated with DMSO as vehicle or 50 μM Mdivi-I or with 5 μM S,S-BMP + 50 μM Mdivi-I for 24 hours. *S,S-*BMP treatment to cells started a passage before the Mdivi-1 treatment and continued till the cells were harvested. To increase yield of lysosomal DNA, cells from five 150 mm x 20 mm plates were pooled as one replicate and processed for lysosome isolation as described. Isolated lysosomes were processed through a genomic DNA extraction kit (ThermoFisher Scientific #K0512) and nucleic acids were eluted in 100 μL water. Isolated DNA was PCR amplified for 60 cycles in technical duplicates and quantified by melting curves as described in the RT-PCR section. Primers used were as follows:

mtATP6-Forward AATCCAAGCCTACGTTTTCACA

mtATP6-Reverse AGTATGAGGAGCGTTATGGAGT

mtCO2-Forward AATCGAGTAGTACTCCCGATTG

mtCO2-Reverse TTCTAGGACGATGGGCATGAAA

mtND1-Forward CTATCACCCTATTAACCACTCA

mtND1-Reverse TTCGCCTGTAATATTGAACGTA

mt-DLoop-Forward CACCCAAGAACAGGGTTTGT

mt-DLoop -Reverse TGGCCATGGGTATGTTGTTAA

---

**Figure S1.**
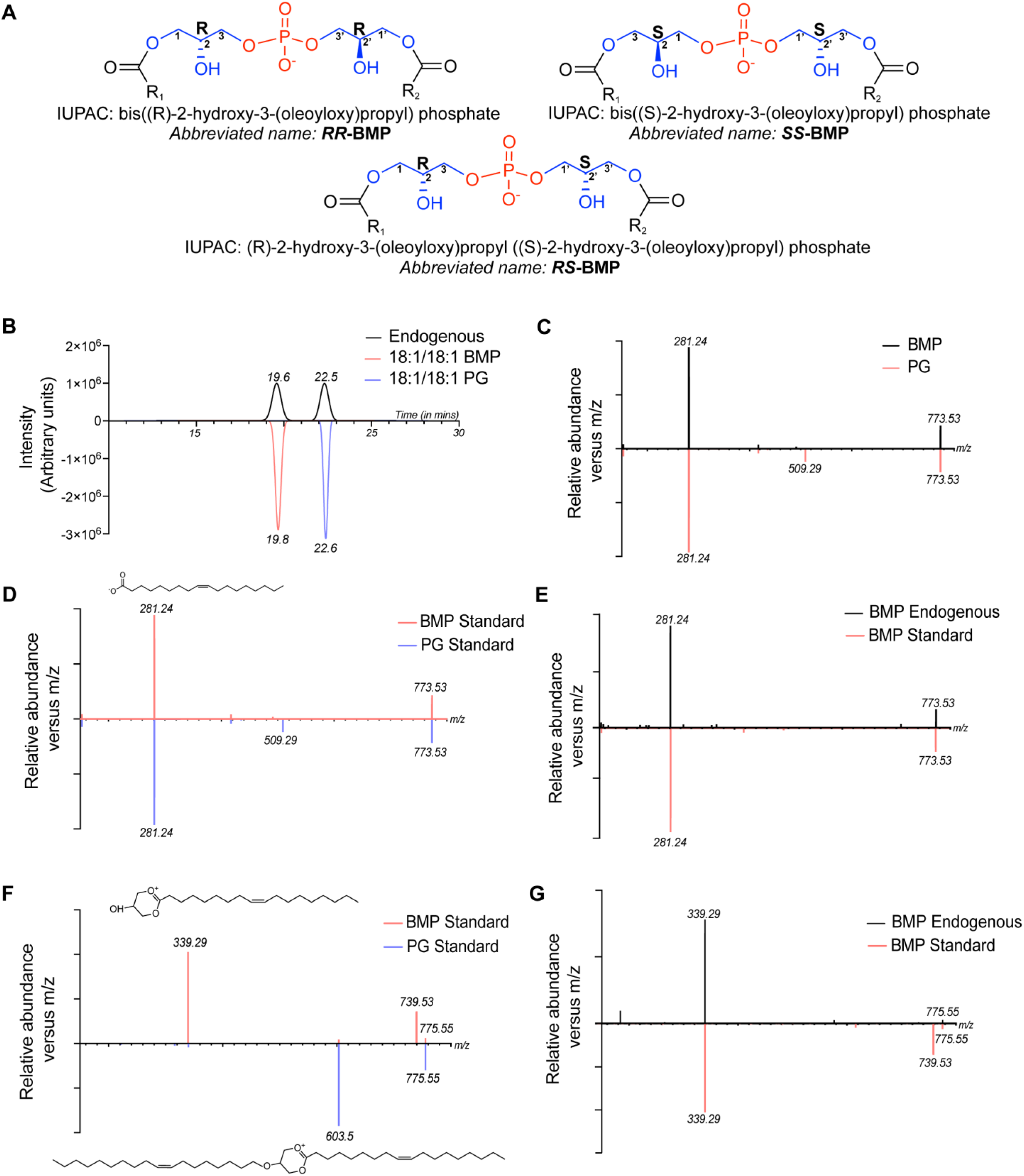
Structure of BMP stereoisomers and LC-MS/MS separation/quantitation method. (A) Structure of BMP stereoisomers. Phosphate in red, glycerol in blue and fatty acids in black. IUPAC name and abbreviated names used for different stereoisomers in the manuscript. (B) Elution trace of 18:1/18:1 PG and 18:1/18:1 BMP on C-30 reverse phase column chromatography. (C) MS/MS fragmentation pattern and alignment of 18:1/18:1 BMP and 18:1/18:1 PG from HEK293T lysates in negative mode of ionization. (D) MS/MS fragmentation pattern and alignment of 18:1/18:1 BMP and 18:1/18:1 PG synthetic standards in negative mode of ionization. (E) MS/MS fragmentation pattern and alignment of 18:1/18:1 endogenous BMP with 18:1/18:1 synthetic standard in negative mode of ionization. (F) MS/MS fragmentation pattern and alignment of synthetic 18:1/18:1 BMP and 18:1/18:1 18:1/18:1 PG in positive mode of ionization. (G) MS/MS fragmentation pattern and alignment of endogenous and synthetic 18:1/18:1 BMP in positive mode of ionization.

**Figure S2.**
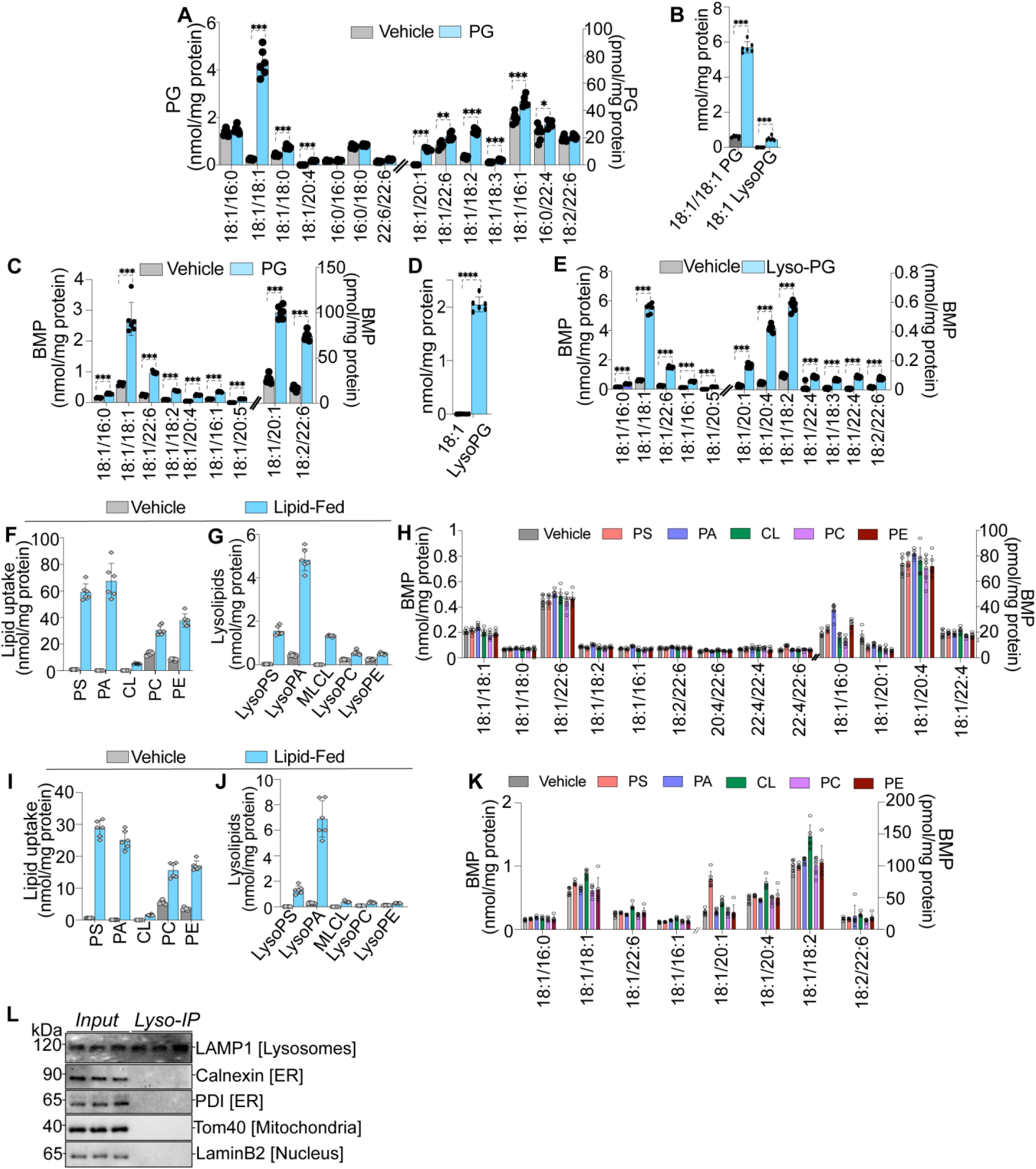
Lyso-PG is an immediate precursor of BMP in mammalian cells. (A) PG species in HMC3 cells after 18:1/18:1 PG incubation for 6 h. (B) 18:1/18:1 PG and 18:1 lyso-PG levels in HEK293T cells after 18:1/18:1 PG incubation for 6 h. (C) BMP species in HEK293T cells after 18:1/18:1 PG incubation for 6 h. (D) 18:1 lyso-PG levels in HEK293T cells after 18:1 lyso-PG incubation for 6 h. (E) BMP species in HEK293T cells after 18:1 lyso-PG incubation for 6 h. (F) Cellular lipids fed to the HMC3 cells for 6 h. (G) Cellular lysophospholipid derivatives of lipids fed to the HMC3 cells for 6 h. (H) BMP in HMC3 cells after lipid incubation for 6 h. (I) Cellular lipids fed to the HEK293T cells for 6 h. (J) Cellular lysophospholipid derivatives of lipids fed to the HEK293T cells after for 6 h. (K) BMP in HEK293T cells after lipid incubation for 6 h. Data presented as mean ± S.D. (n=6 per group). \**p < 0.05*; \*\**p < 0.01*; \*\*\**p < 0.001*; \*\*\*\**p <0.0001* vehicle versus lipid feeding (multiple t-test). (L) Representative western blot showing rapid isolation of lysosomes from HMC3 cells using TMEM192-3xHA tag-mediated immuno-isolation.

**Figure S3.**
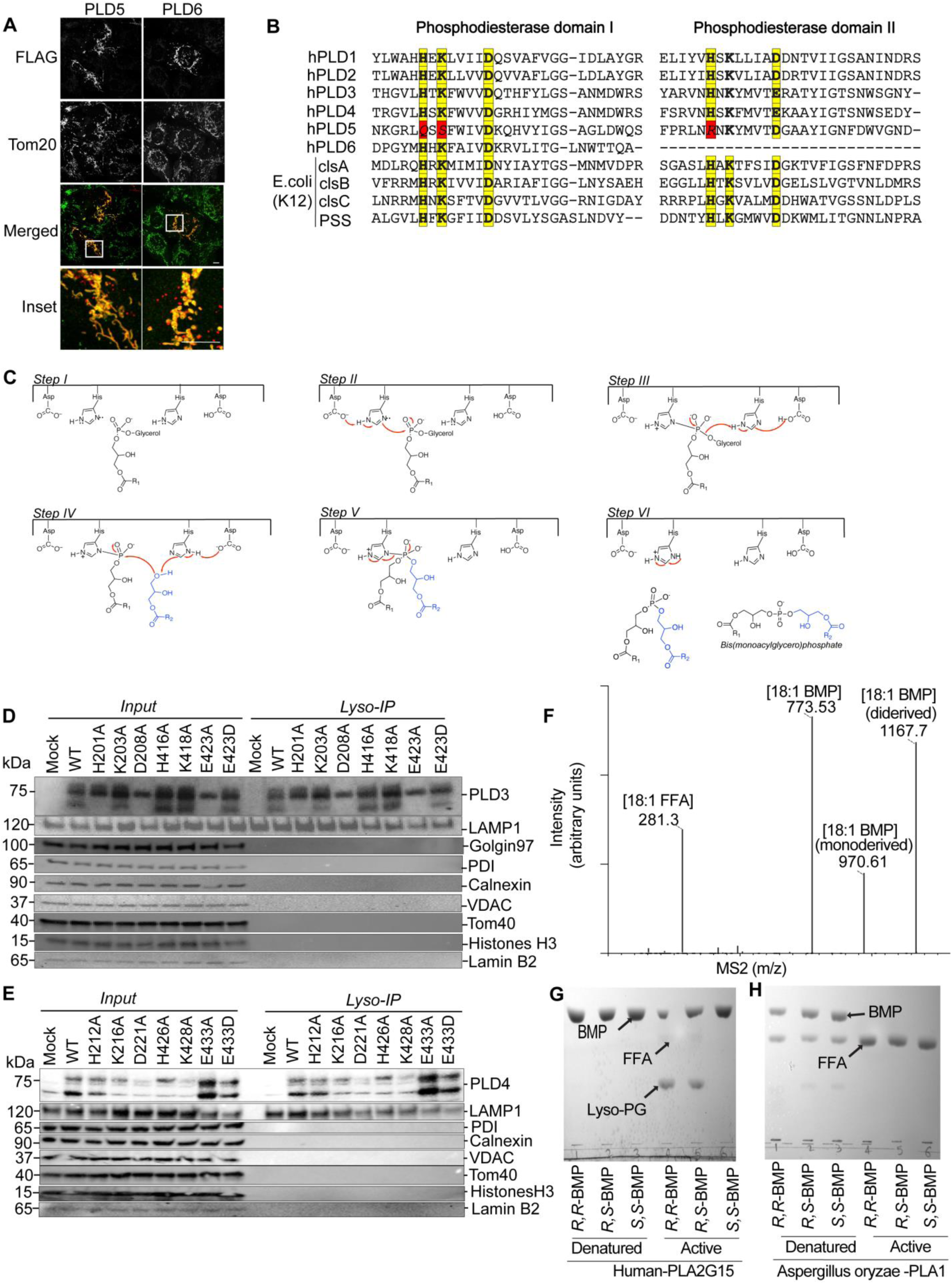
PLD enzymes and subcellular localization of mutants. (A) Confocal microscopy images probing subcellular localization of PLD5 and PLD6 in the HMC3 cells, PLD5-6 shown in red, Tom20 shown in green. Scale bar =10 *μm*. (B) Conserved phosphodiesterase domains HxKxxxxD of PLDs. (C) Proposed arrow pushing mechanism involving two catalytic triads of PLDs for transphosphatidylation reaction and BMP synthesis. Lysines of the catalytic triad that presumably stabilize phosphate group of lipids are removed for simplicity. (D) Western blot probing expression of catalytically impaired mutants of PLD3 in lysosomes isolated from HEK293T cells. (E) Western blot probing expression of catalytically impaired mutants of PLD4 in lysosomes isolated from HEK293T cells. (F) Fragment spectra of BMP derivatized with two mol-equivalents of (R)-(-)-1-(1-naphthyl)ethyl isocyanate (R-NIC). m/z=1167.7 corresponds to 18:1/18:1 BMP derivatized with two molecules of R-NIC, m/z=970.61= 18:1/18:1 BMP derivatized with one molecule of R-NIC, m/z=773.53 corresponds to 18:1/18:1 BMP after both R-NIC molecules fall off, m/z=281.3 corresponds to 18:1 free fatty acid. (G) Image of TLC plate demonstrating human PLA2G15 activity on different BMP stereoisomers. PLA2G15 cleaves R,R-BMP and R,S-BMP but not S,S-BMP. (H) Image of TLC plate showing robust and indiscriminate hydrolysis of all BMP stereoisomers by *Aspergillus oryzae* PLA1 (Sigma-Aldrich #L3295).

**Figure S4.**
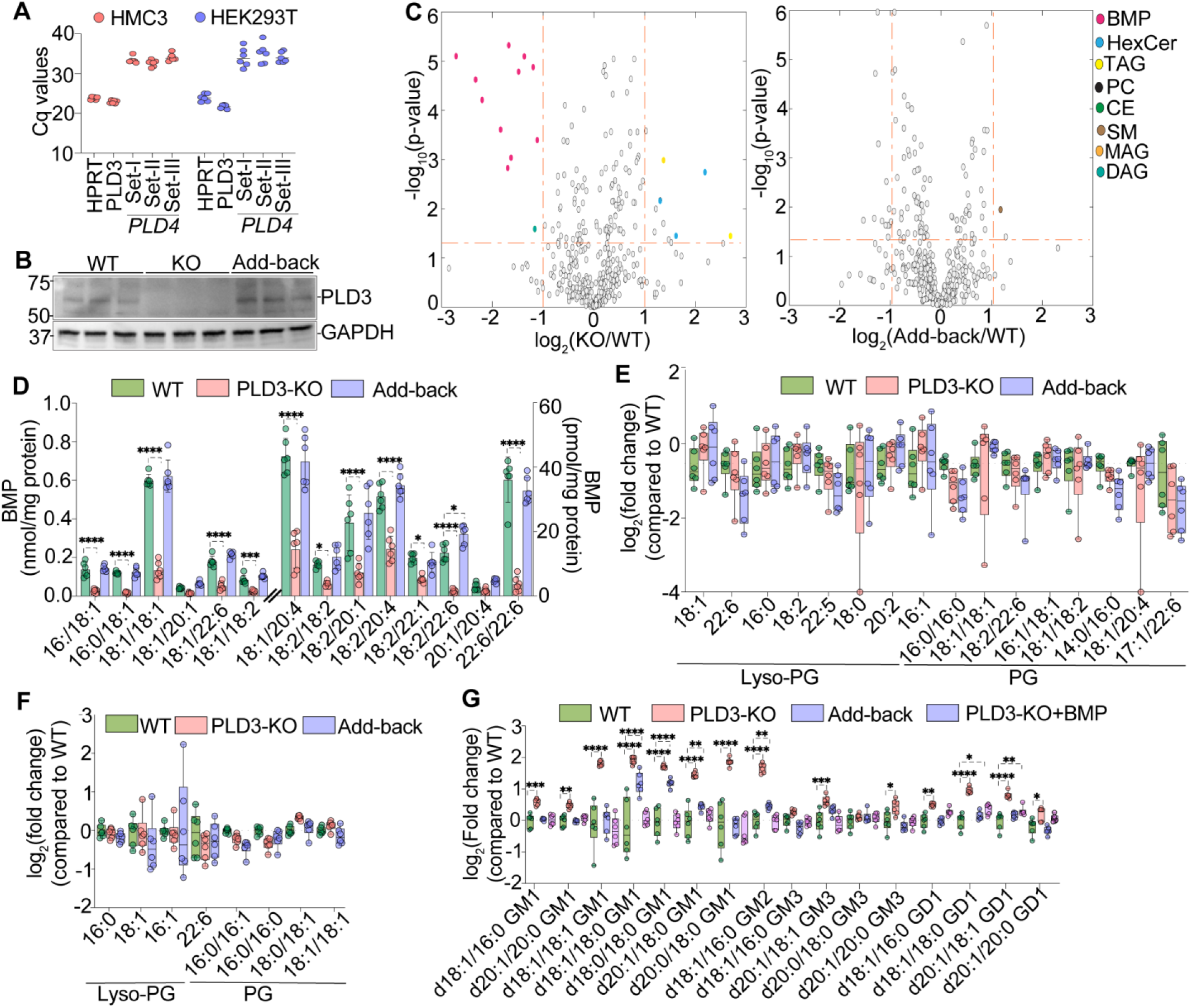
Reduced BMP levels in PLD3-deficient mammalian cells. (A) RT-PCR Cq values for PLD3 and PLD4 in HMC3 and HEK293T cells. (B) Western blot probing presence of PLD3 in WT, PLD3 KO and add-back (i.e., PLD3 KO rescued by stable expression of PLD3) HEK293T cells. (C) Volcano plots of whole cell lipid measurement of PLD3 KO plotted against WT and PLD3 add-back plotted against WT HEK293T cells with log_2_-fold changes (ratio of relative abundance, x-axis) and log_10_ *p*-values (y-axis). BMPs were most decreased depicted as red dots. *p-*values calculated by two-sample t-test. (D) BMP species in WT, PLD3 KO and PLD3 add-back HEK293T cells. (E) Relative changes in lyso-PG and PG species in WT, PLD3 KO and PLD3 add-back HMC3 cells. (F) Relative changes in lyso-PG and PG species in WT, PLD3 KO and PLD3 add-back HEK293T cells. (G) Ganglioside species in WT, PLD3 KO, PLD3 add-back or PLD3 KO cells fed with *S,S*-BMP in HEK293T cells. Data presented as mean ± S.D. (n=6 per group). \**p < 0.05*; \*\**p < 0.01*; \*\*\**p < 0.001*; \*\*\*\**p <0.0001* (two-way ANOVA multiple comparison with Dunnett’s correction).

**Figure S5.**
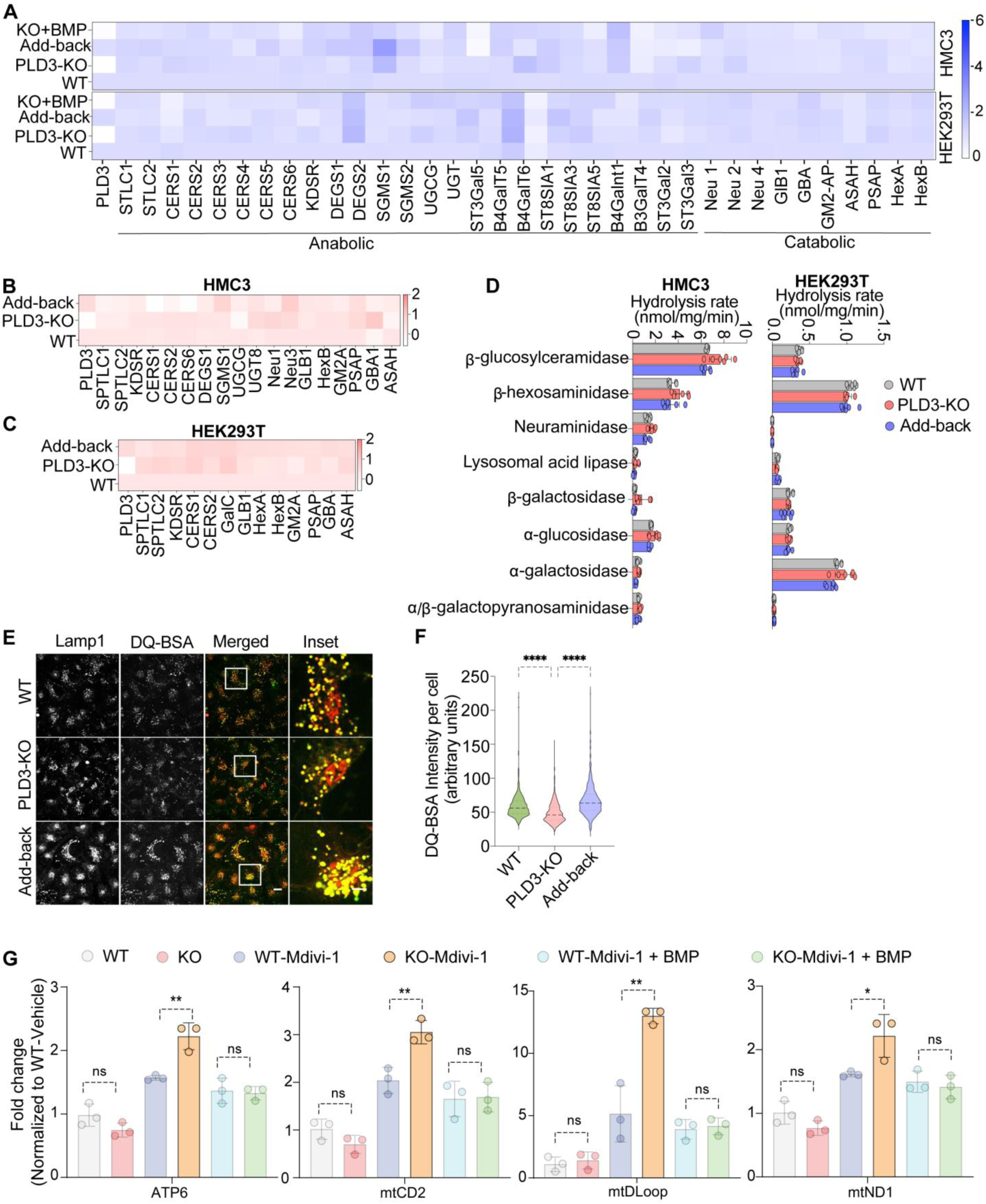
PLD3 deficiency results in lysosomal dysfunction without change in abundance of ganglioside metabolism enzyme. (A) Heat map analysis of relative transcript levels of ganglioside metabolic genes in the PLD3 KO, PLD3 add-back and PLD3 KO cells fed with *S,S*-BMP against WT HMC3 and HEK293T cells. (B) Heat map analysis of the relative abundance of glycosphingolipid metabolism proteins from PLD3 KO, add-back against WT HMC3 cells. (C) Heat map analysis of the relative abundance of glycosphingolipid metabolism proteins from PLD3 KO, PLD3 add-back against WT HEK293T cells; data plotted as mean of the log_2_ (fold-changes). (D) Sugar lipid catabolic activities using artificial substrate mimetics in extracts of HMC3 and HEK293T cells. (E) Confocal microscopy images of cleaved DQ-BSA (red) and LysoTracker (green) demonstrating decreased DQ-BSA signal in the lysosomes of PLD3 KO HMC3 cells, compared to WT and PLD3 add-back cells. Scale bar =10 *μm*. (F) Quantification of DQ-BSA signal (green). \*\*\*\**p < 0.0001* (two-way ANOVA multiple comparison with Dunnett’s correction). (G) Quantification of mitochondrial DNA by PCR in the lysosomes of WT and PLD3 knock out HMC3 cells treated with DMSO, Mdivi-1 (mitophagy inducer) and *S,S*-BMP+Mdivi-1. Data presented as mean ± S.D. (n=3 per group). \**p<0.05*, \*\**p<0.001* (two-tailed unpaired t-test).

**Figure S6.**
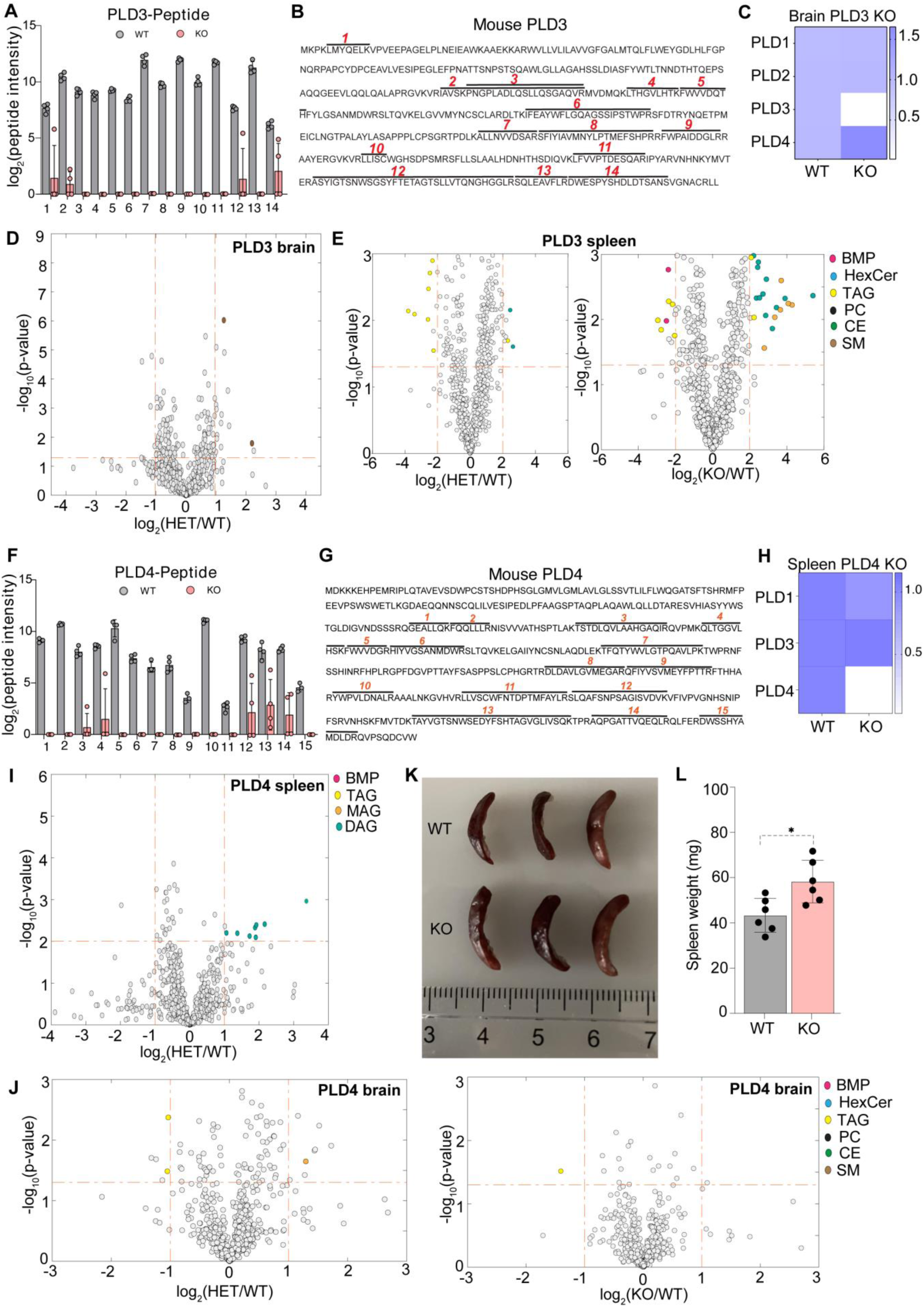
Deficiency of PLD3 or PLD4 alters BMP levels in murine tissues. (A) Mass spectrometry data demonstrating loss of PLD3 peptides in brain of PLD3 KO mice. (B) Map of PLD3 peptides detected in mass spectrometry on full length mouse PLD3 protein sequence. (C) Heat map demonstrating levels of PLD1-4 in brain of PLD3 KO mice. Minor increase in PLD4 levels in brain of PLD3 KO mice. Data plotted as fold change normalized to levels in WT mice. (D) Volcano plot of lipid measurements in the brain of PLD3 heterozygous (HET) versus PLD3 WT mice with log_2_-fold changes (ratio of relative abundance, x-axis) and log_10_ *p-*values (y-axis). (E) Volcano plot of lipid measurements in the spleen of heterozygous PLD3 +/- (HET) versus PLD3 WT mice and PLD3 KO versus PLD3 WT with log_2_-fold-changes (ratio of relative abundance, x-axis) and log_10_ *p-*values (y-axis). (F) Mass spectrometry data demonstrating loss of PLD4 peptides in spleen of PLD4 KO mice. (G) Map of PLD4 peptides detected in mass spectrometry on full length mouse PLD4 protein sequence. (H) Heat map demonstrating levels of PLD1,3,4 in spleen of PLD4 KO mice. Data plotted as fold change normalized to levels in WT mice. (I) Volcano plot representation of lipid measurements in the spleen of heterozygous PLD4 +/- (HET) versus PLD4 WT mice with log_2_-fold changes (ratio of relative abundance, x-axis) and log_10_ *p-*values (y-axis). (J) Volcano plot representation of lipid measurements in the brain of PLD4 HET versus PLD4 WT mice and PLD4 KO versus PLD4 WT with log_2_-fold change (ratio of relative abundance, x-axis) and log_10_ *p-*value (y-axis). *p-*values calculated by two-sample t-test. (K) Representative image demonstrating splenomegaly in PLD4 KO mice, compared to PLD4 WT mice. (L) Wet tissue weight of spleen from 3–4 months old PLD4 KO and WT mice. *p-*Values calculated using Student’s t-test.

**Figure S7.**
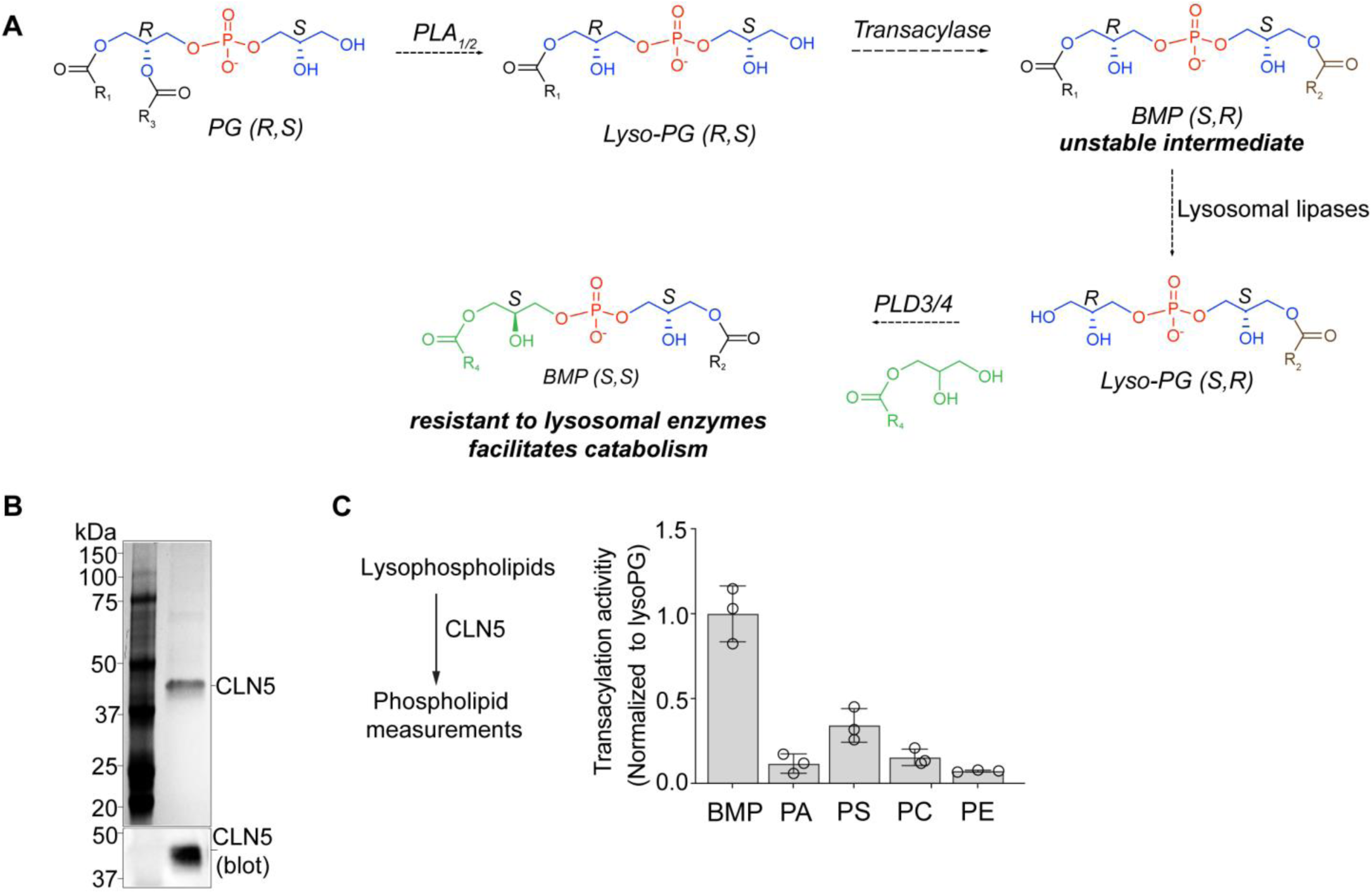
Recombinant CLN5 synthesizes R,racemic-sBMP *in vitro*. (A) Proposed *S,S-*BMP synthesis pathway involving transacylation and transphosphatidylation reactions in mammals. (B) Coomassie-stained SDS-PAGE gel of purified recombinant human CLN5. (C) Transacylation activity of recombinant CLN5. All lysophospholipids used as substrate contained 18:1 free fatty acid as acylated chain.

